# Devaluation of response-produced safety signals reveals circuits for goal-directed versus habitual avoidance in dorsal striatum

**DOI:** 10.1101/2024.02.07.579321

**Authors:** Robert M. Sears, Erika C. Andrade, Shanna B. Samels, Lindsay C. Laughlin, Danielle M. Moloney, Donald A. Wilson, Matthew R. Alwood, Justin M. Moscarello, Christopher K. Cain

## Abstract

Active avoidance responses (ARs) are instrumental behaviors that prevent harm. Adaptive ARs may contribute to active coping, whereas maladaptive avoidance habits are implicated in anxiety and obsessive-compulsive disorders. The AR learning mechanism has remained elusive, as successful avoidance trials produce no obvious reinforcer. We used a novel outcome-devaluation procedure in rats to show that ARs are positively reinforced by response-produced feedback (FB) cues that develop into safety signals during training. Males were sensitive to FB-devaluation after moderate training, but not overtraining, consistent with a transition from goal-directed to habitual avoidance. Using chemogenetics and FB-devaluation, we also show that goal-directed vs. habitual ARs depend on dorsomedial vs. dorsolateral striatum, suggesting a significant overlap between the mechanisms of avoidance and rewarded instrumental behavior. Females were insensitive to FB-devaluation due to a remarkable context-dependence of counterconditioning. However, degrading the AR-FB contingency suggests that both sexes rely on safety signals to perform goal-directed ARs.

---

Active avoidance (AA) is a form of response learning that allows subjects to suppress fear and neutralize threats. AA likely evolved to expand the defensive behavior repertoire beyond a narrow set of innate, inflexible reactions (i.e. fight, flight, freezing). Avoidance response (AR) learning may even underlie complex defensive behaviors that could not have evolved under predatory pressure. For instance, a commuter may buckle their seatbelt to protect against future injury or slam their brakes to prevent imminent harm. Clinically, AA is important because adaptive ARs may contribute to active coping and resilience^1^. Conversely, inappropriate or persistent avoidance habits are thought to protect irrational fears from extinction^2^ and contribute to disorders of control like OCD^3^.

AA is typically studied by presenting subjects with a warning stimulus (WS; e.g., sound) that predicts the delivery of an aversive unconditioned stimulus (US; e.g., shock). This transforms the WS into a conditioned threat that arouses fear and triggers reactions like freezing. Subjects eventually emit the AR (e.g., barpress) and gain exposure to avoidance contingencies. As training continues, fear reactions to the WS are progressively replaced by ARs.

Progress in understanding the associative structures and neurobiological mechanisms of AA was stalled by theoretical disputes about the reinforcement mechanism and misconceptions about the mediating role of fear^4,5^. Prominent theories focused on stimulus-response (S-R) learning via negative reinforcement (i.e. fear-reduction at WS-termination vs. US-omission)^6,7^. AA research also failed to fully capitalize on modern conceptions of instrumental behavior developed using appetitive reinforcers^8,9^. This work moved beyond the notion that instrumental training produces only reflexive S-R habits. Subjects also learn goal-directed responses that depend on the action-outcome (A-O) contingency and outcome value. Goal-directed responding is typically diagnosed with outcome-devaluation or contingency degradation tests^8,9^. Thus, barpressing for food is goal-directed if reducing the food’s value, via satiation or counterconditioning, reduces barpressing. The same can be said if food delivered non-contingently (independent of behavior) reduces barpressing. S-R habits are usually inferred by insensitivity to these tests. However, the situation is more complicated for AA as there is little agreement about what reinforces ARs. If WS-termination and/or US-omission contribute, it is unclear how one could verifiably devalue these outcomes, or deliver more of them. Identifying a clear reinforcement mechanism would establish that AA is instrumental and help differentiate between goal-directed versus habitual ARs.

A less studied hypothesis is that ARs are positively reinforced by response-produced safety signals that are generated via their inverse relationship with the WS and US^7,10–12^. Explicit feedback (FB) cues that immediately follow ARs develop conditioned inhibition (safety) and reinforcing properties^10,13–16^. It is unknown, however, if safety signals behave as valued outcomes supporting goal-directed avoidance, akin to food delivery in a barpressing experiment. To our knowledge, no AA studies have attempted to devalue response-produced safety signals. To examine this, we designed a devaluation protocol where AR-FB cues are counterconditioned by pairing with shock. We also present data on a novel contingency degradation test, where non-contingent FB cues are presented during avoidance.

If ARs are positively reinforced by safety, this raises the exciting possibility that AA relies on instrumental brain circuits already identified by appetitive studies. Moderately trained instrumental responses are sensitive to outcome devaluation and are suppressed by manipulations that impair neural activity in posterior dorsomedial striatum (pDMS)^17,18^. In contrast, instrumental responses that are overtrained or trained on interval schedules are insensitive to outcome devaluation and are suppressed by manipulations that impair dorsolateral striatum (DLS)^18,19^. Importantly, however, this latter effect is only evident in subjects that also received outcome devaluation or contingency degradation. This is because goal-directed responding mediated by a parallel pDMS circuit regains control when DLS-mediated habits are impaired^19^. Thus, if goal-directed and habitual avoidance memories co-exist in different brain circuits, a tool like outcome devaluation may be necessary to identify and study these circuits. Impairing either circuit alone may produce no change in overt avoidance behavior.

Here, we show that counterconditioning of response-produced safety signals impairs moderately trained, but not overtrained, ARs in male rats. We also demonstrate that moderately trained ARs depend on pDMS, and overtrained ARs depend on DLS, as in appetitive studies. Further, we found that ARs in female rats are insensitive to devaluation due to a remarkable context dependence of counterconditioning. Lastly, we show contingency degradation data suggesting that both sexes rely on safety signals for goal-directed avoidance.

## Results

### Avoidance is acquired similarly in male and female rats

Overtraining is frequently used to induce habitual responding in studies where responses are rewarded^20^. To study the development of avoidance habits, we used a signaled active avoidance (SigAA) shuttlebox procedure in rats and varied the amount of training. Fig. 1 shows SigAA training data from multiple behavioral experiments. The percentage of good avoiders was similar in male and female rats (Fig. 1b). Good avoiders approached 100% successful avoidance by the fifth training session and average AR latencies settled around 5s after WS onset (Fig. 1c,d). Interestingly, variation in AR latencies declined with overtraining even as average AR latencies remained constant (Fig. 1d, right). This invariant response topography is a positive sign of automaticity^21^, and possibly habitual control^22^. Nearly identical patterns of AR learning and performance were observed for males and females in both training cohorts. The only sex differences observed were slightly more shuttles during the acclimation and inter-trial intervals (ITIs) for females (Extended Fig. 1).

**Fig. 1.**
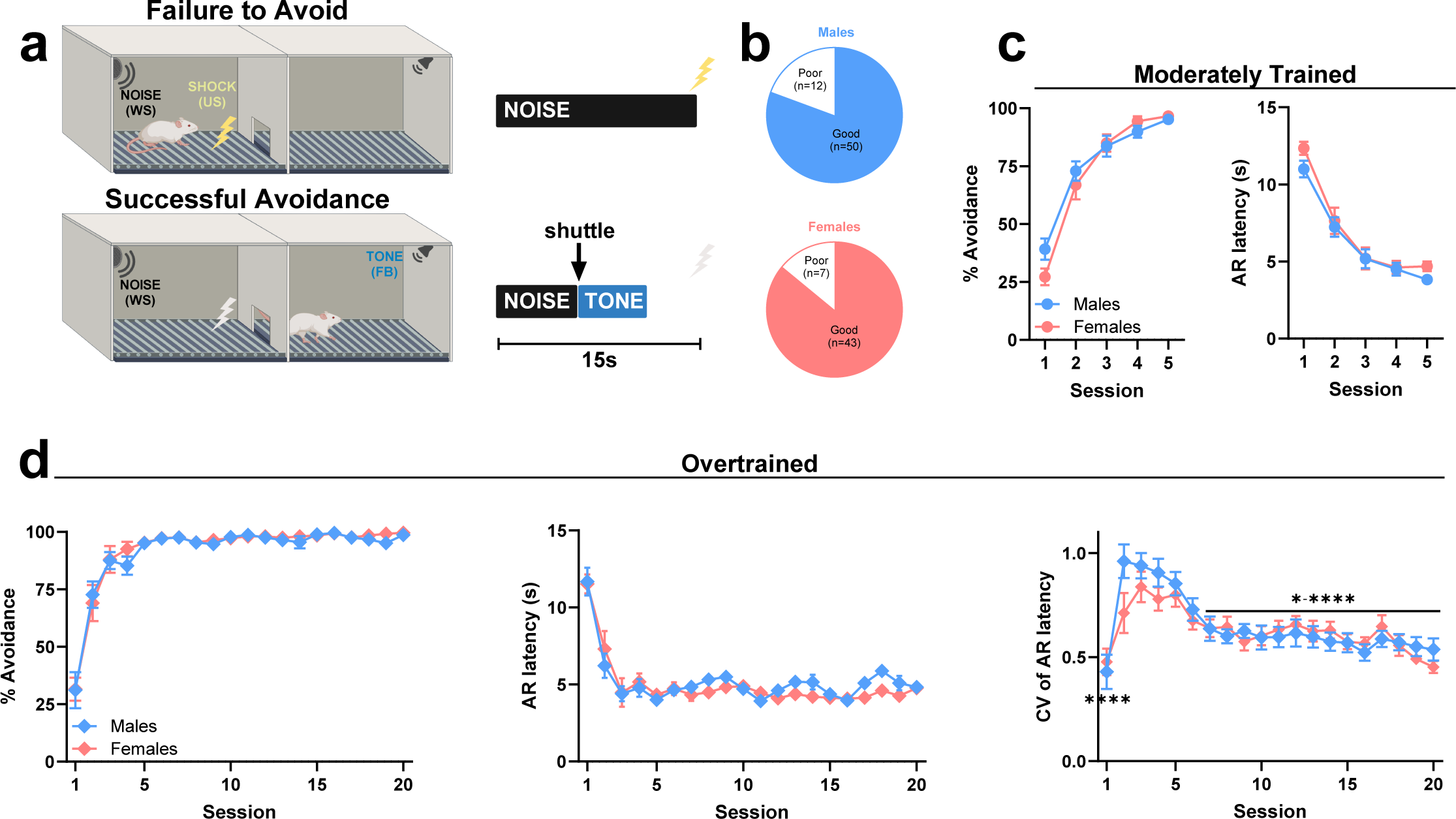
Acquisition of SigAA behavior in male and female rats. **a**, Schematic of the shuttlebox SigAA paradigm. **b**, Proportion of good versus poor avoiders in males and females from multiple behavioral experiments. **c**, Avoidance behavior during moderate training in male (n=35) and female (n=26) rats. Two-way ANOVA. % Avoidance: F_4,236_=2.28. AR latency: F_4,236_= 0.73. **d**, Avoidance behavior during overtraining in male (n=15) and female (n=17) rats. Two-way ANOVA. % Avoidance: F_19,570_=0.34. AR latency: F_19,570_=0.995. For each animal, a significant large-sized session effect (F = 8.37, p < 0.001, η^2^ = 0.17) was apparent for AR latency. We therefore calculated coefficients of variation (CV) to assess the extent of variability across sessions using Two-way ANOVA followed by Tukey’s multiple comparisons test. Here, CV is inversely related to stereotyped behavior. CV of AR latency: F_19,570_=1.24 (Session x Sex), F_19,570_=11.89 (Session); Session comparisons, Session 5 versus Session 1 ****p<0.0001, Session 5 versus Sessions 7-20 *p=0.02-****p<0.0001. All line graphs are shown as mean ± s.e.m.

### Devaluation of response-produced safety signals impairs moderately trained avoidance in male rats

To test whether goal-directed avoidance depends on safety signals, male rats received moderate SigAA training followed by paired (FB-Devalued and FB-Control groups) or explicitly unpaired (FB-Valued group) counterconditioning in a separate chamber (Fig. 2a). The unpaired procedure controlled for stimulus exposure during counterconditioning but preserved the safety value of the tone. The FB-Control group was treated identically to FB-Devalued group except the FB stimulus used during SigAA training was different (house light off). This allowed us to assess *retardation of acquisition*^23^ for FB-Devalued rats during counterconditioning compared to FB-Control rats that also had SigAA training, but the tone was novel. It also allowed us to assess whether any AR impairment induced by counterconditioning was specific to the FB stimulus used during SigAA training. After counterconditioning, rats were returned to the shuttleboxes for two devaluation tests (15 presentations of the WS). Shuttling had no effect on WS duration, and no FB or shocks were presented. After this, a final Tone Fear Test was conducted in a novel open field to confirm that counterconditioning was effective (FB-Valued vs. FB-Devalued) and to assess retardation of acquisition (FB-Devalued vs. FB-Control) in a neutral context.

**Fig. 2.**
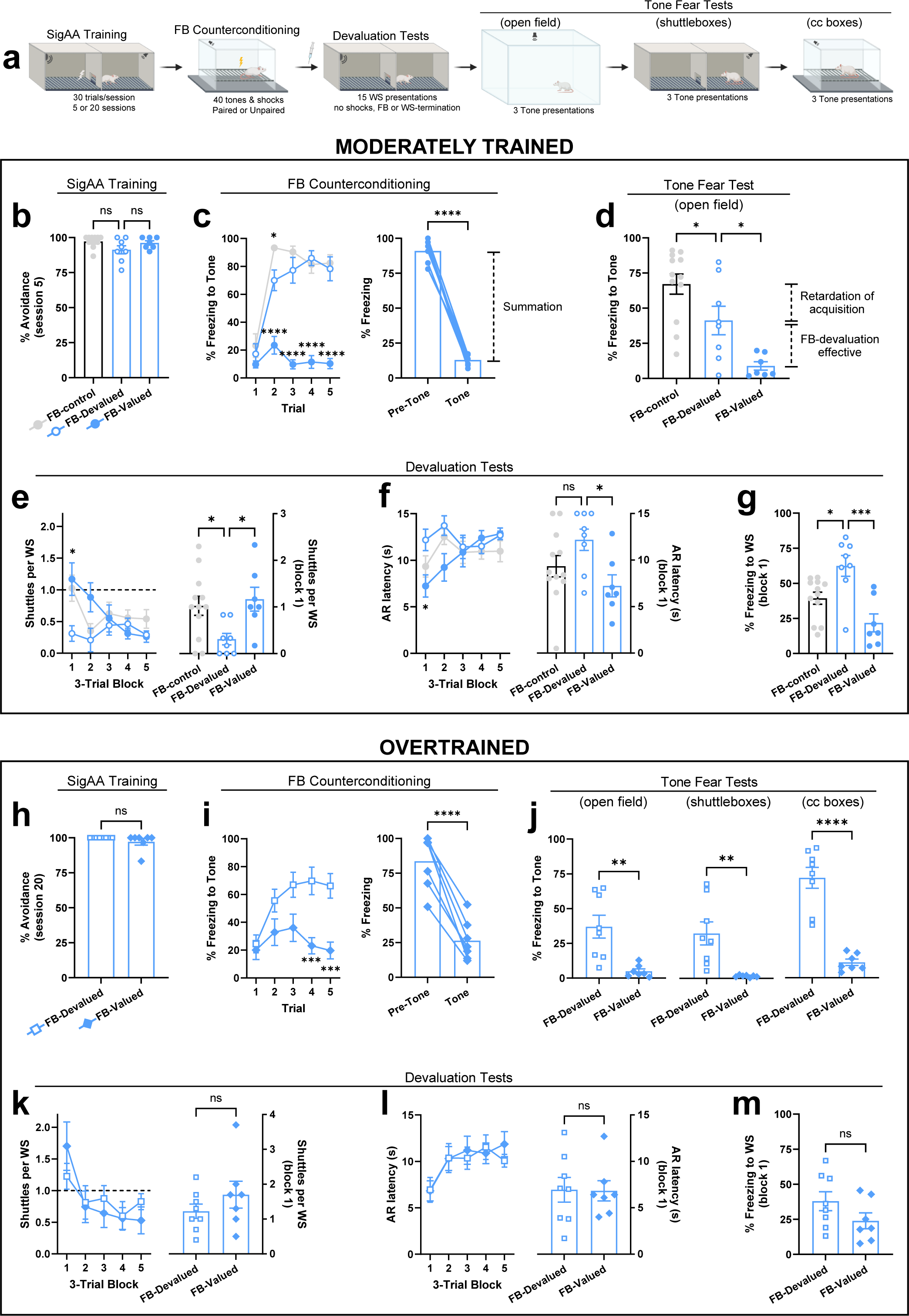
Devaluation of response-produced safety signals after moderate training versus overtraining in male rats. **a**, Schematic of the devaluation paradigm. Panels **b-g**: moderately trained rats (FB-Control: n=12, FB-Devalued: n=8, FB-Valued: n=7). Panels **h-m**: overtrained rats (FB-Devalued: n=8, FB-Valued: 7). Note that moderately trained rats received only one Tone Fear Test in the open field. Additional Tone Fear Tests in the counterconditioning boxes and shuttleboxes were later added for overtrained rats to address differences between males and females. **b**, SigAA performance during session 5 of training. One-way ANOVA. % Avoidance: F_2,24_=2.938. **c**, Left, Tone freezing during the first 5 trials of counterconditioning. Two-way ANOVA followed by Dunnett’s test: F_8,120_=5.95. Group comparisons, FB-Devalued versus FB-Control, Trial 2 *p= 0.014; FB-Devalued versus FB-Valued, Trials 2-5 ****p<0.0001. Right, Pre-Tone versus Tone freezing during first 5 trials of counterconditioning for FB-Valued rats. Paired t-test: t_6_=21.68 ****p<0.0001. **d**, Tone freezing in the novel open field. One-way ANOVA followed by Dunnett’s test: F_2,24_=13.95. Group comparisons, FB-control versus FB-Devalued *p=0.04, FB-Devalued versus FB-Valued *p=0.02. **e**, Left, shuttles per WS for both Devaluation Tests. Two-way ANOVA followed by Dunnett’s test: F_8,96_=4.41. Group comparisons, FB-Devalued versus FB-control: Trial 1 *p=0.02; FB-Devalued versus FB-Valued: Trial 1 *p=0.03. Right, shuttles per WS for Block 1 of Devaluation Tests. One-way ANOVA followed by Dunnett’s test: F_2,24_=4.445. Group comparisons, FB-Devalued versus FB-Control *p=0.03, FB-Devalued versus FB-Valued *p=0.02. **f**, Left, latency to first AR for both Devaluation Tests. Two-way ANOVA followed by test: F_8,96_=3.05. Group comparisons, FB-Devalued versus FB-Valued, Block 1 *p=0.02. Right, latency to first AR during Block 1 of Devaluation Tests. One-way ANOVA followed by Dunnett’s test: F_2,24_=3.67. Group comparisons, FB-Devalued versus FB-Control p=0.16, FB-Devalued versus FB-Valued *p=0.02. **g**, WS freezing for Block 1 of Devaluation Tests. One-way ANOVA followed by Dunnett’s test: F_2,24_=10.42. Group comparisons, FB-Devalued versus FB-Control *p=0.01, FB-Devalued versus FB-Valued ***p=0.0003. **h**, SigAA performance during session 20 of overtraining for rats assigned to FB-Devalued versus FB-Valued groups. Unpaired t-test: t_13_=1.31 p=0.21. **i**, Left, Tone freezing during the first 5 trials of counterconditioning. Two-way ANOVA followed by Šídák’s test: F_4,52_=4.59. Group comparisons, FB-Valued versus FB-Devalued, Trial 4 ***p=0.0009, Trial 5 ***p=0.001. Right, Pre-Tone versus Tone freezing during first 5 trials of counterconditioning for FB-Valued rats. Paired t-test: t_6_=9.71 ***p<0.0001. **j**, Tone freezing in the novel open field (left), shuttleboxes (middle), and counterconditioning boxes (right). Unpaired t-tests: t_13_=3.58 **p=0.003 (left); t_13_=3.44 **p=0.004 (middle); t_13_=7.27 ****p<0.0001 (right). **k**, Left, shuttles per WS for both Devaluation Tests. Two-way ANOVA: F_4,52_=2.07. Right, shuttles per WS for Block 1 of Devaluation Tests. Unpaired t-test: t_13_=1.14 p=0.28. **l**, Left, latency to first AR for both Devaluation Tests. Two-way ANOVA: F_4,52_=0.81. Right, latency to first AR during Block 1 of Devaluation Tests. Unpaired t-test: t_13_=0.06 p=0.95. **m**, WS freezing for Block 1 of Devaluation Tests. Unpaired t-test: t_13_=1.56 p=0.14. Data in **c** (right) and **i** (right) are shown as mean + individuals. All other data in the figure are shown as mean ± s.e.m.

Rats in all moderately trained groups displayed high levels of avoidance responding prior to FB devaluation (Fig. 2b). Freezing measured during the first five trials of counterconditioning revealed several important group differences (Fig. 2c). First, tone-freezing increased quickly in the FB-Devalued group but remained low in the FB-Valued group, indicating that paired counterconditioning had the intended effect. Second, tone-freezing in the FB-Control group on Trial 2 was higher than FB-Devalued rats, consistent with retardation of acquisition for FB-Devalued rats that began the session treating the tone as a safety signal. Third, freezing in the FB-Valued group was high before tone presentations, due to context fear induced by unpaired shocks, but ceased during tone presentations (Fig. 2c, right). This dramatic *summation* effect^23^ is another indicator that tones used as FB during SigAA training become strong safety signals. These conclusions were supported by freezing behavior during the Tone Fear Test in the open field (Fig. 2d). FB-Devalued rats showed tone-freezing intermediate to high-freezing FB-Control rats and low-freezing FB-Valued rats. Behaviors during the Devaluation Tests indicate that FB counterconditioning significantly impaired avoidance, especially during the first testing block, when responding is most clearly related to a memory of the instrumental outcome^24^. Shuttling during the WS was much lower for FB-Devalued rats compared to both control groups (Fig. 2e). Consistent with this, AR latencies were significantly longer for FB-Devalued rats (Fig. 2f). Finally, we also measured freezing to the WS during the first block of testing. Freezing is a fear-related behavior that competes with anxiety-related avoidance^4^, and multiple studies have found that suppressing AA disinhibits freezing^25–27^. This was also true in our experiment as FB-Devalued rats froze significantly more than both control groups during WS presentations (Fig. 2g). Together, these data indicate that avoidance FB stimuli become safety signals during SigAA training, these safety signals are devalued by counterconditioning, and this devaluation impairs moderately trained avoidance even when the devalued FB stimulus is not presented during the test. This is consistent with goal-directed responding where the decision to avoid depends on a memory of the current outcome value. Importantly, data from the FB-Control group also show that our devaluation effect was specific; strong tone-shock conditioning had no effect on ARs trained with a different FB stimulus.

We next conducted the same analyses in overtrained male rats. Additional Tone Fear Tests were conducted in the shuttleboxes and counterconditioning boxes to address sex differences (discussed later). We again saw evidence for conditioned inhibition to the tone (summation; Fig. 2i, right). We also confirmed that counterconditioning was effective and transferred to all contexts tested (Fig. 2j). However, FB counterconditioning had no effect on overtrained avoidance during the Devaluation Tests (Fig. 2k-m). These data indicate that FB counterconditioning remains effective after overtraining, but ARs are insensitive to devaluation, most likely because they are mediated by S-R associations that bypass outcome value when controlling behavior.

### pDMS is required for goal-directed active avoidance

To test the role of pDMS, aDMS, and DLS in moderately trained SigAA we used a viral κ-opioid receptor DREADD (KORD) strategy to reversibly suppress neuronal activity in dorsal striatum. KORD is an engineered Gαi-coupled receptor with no natural ligand that hyperpolarizes neurons when bound to Salvinorin B (SalB), an otherwise inert compound^28,[1]^. We first expressed KORD unilaterally in dorsal striatum neurons (see Methods). We then monitored field activity in awake rats via bilateral chronically implanted electrodes before and after SalB (i.v., 2.5 mg/kg; Extended Fig. 3a). Theta band oscillations are a primary LFP signature of activity in dorsal striatum^29,30^, and neurons in dorsal striatum become entrained to theta-band frequencies during instrumental learning^29^. Based on assessment of the full mean FFT spectrum, the largest effect of SalB was suppression of theta (9-15Hz) in the KORD hemisphere (Extended Fig. 3c). Spontaneous theta was significantly depressed compared to baseline and compared to the control hemisphere (Extended Fig. 3d), which was not significantly affected by SalB. Together, these data confirm that KORD/SalB selectively suppresses behaviorally relevant neural activity in dorsal striatum.

We next expressed KORD bilaterally in pDMS of male rats before moderate SigAA training (Fig. 3a,b). Each rat then received two AR test sessions after Vehicle or SalB injections (counterbalanced). AR tests consisted of 15 presentations of the WS where shuttling had no effect on WS duration, and no FB stimuli or shocks were presented. Note that we confirmed high test-retest reliability before pursuing this within-subjects strategy with KORD (Extended Fig. 2). At the end of SigAA training, rats assigned to receive Vehicle-first or SalB-first displayed equivalent avoidance (Fig. 3c). Compared to Vehicle, SalB significantly reduced shuttling during the WS, especially during the first testing block (Fig. 3d). There was also a trend towards longer AR latencies during the SalB test (Fig. 3e). Consistent with an impairment of avoidance, freezing to the WS was increased by SalB (Fig. 3f). In contrast, SalB had no effect on moderately trained avoidance or WS-freezing in rats expressing KORD in aDMS (Extended Fig. 3f-j) or DLS (Fig. 3g-k). Further, SalB also had no effect in overtrained rats expressing KORD bilaterally in pDMS (Extended Fig. 3k-o), presumably because S-R associations stored outside pDMS maintain responding. This aligns with studies showing that pDMS lesions fail to impair habitual responding trained with food rewards^19^. Together, these findings indicate that pDMS is critical for performance of goal-directed, but not habitual, ARs. They also suggest that DMS-mediated control of goal-directed ARs occurs before the development of DLS-mediated habits (Fig. 3l-n). Note that KORD/SalB occasionally reduced shuttles during the acclimation or ITI periods of extinction tests, suggesting a modest influence on overall activity (Extended Figs. 3,4). However, these effects were inconsistent and can’t explain the patterns of responding observed during the WS.

**Fig. 3.**
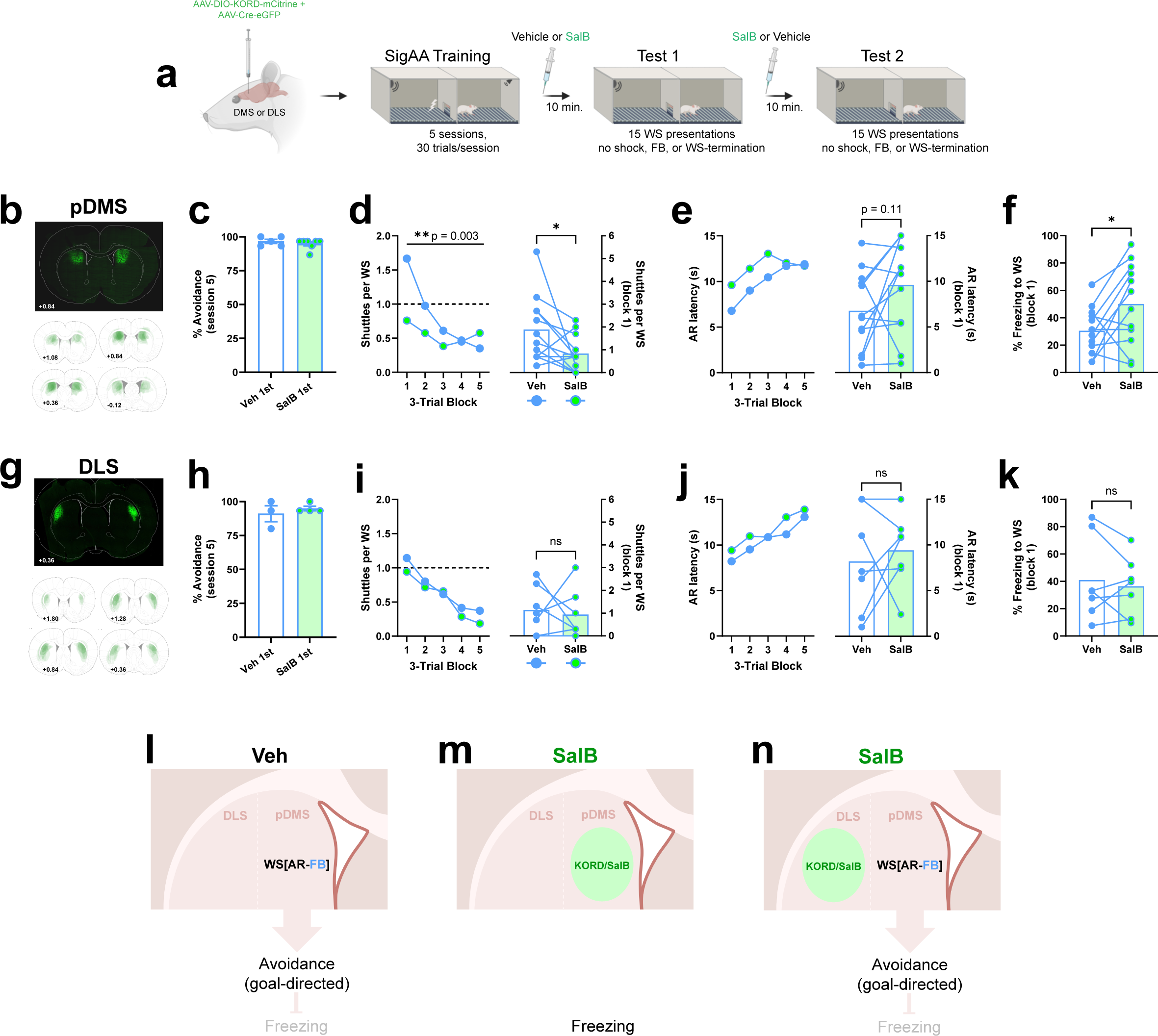
Inhibition of pDMS impairs goal-directed avoidance in male rats. **a**, Schematic of the experimental paradigm. Panels **b-f**: rats expressing KORD in pDMS (n=12). Panels **g-k**: rats expressing KORD in DLS (n=7). **b**, Top, example of bilateral KORD expression in pDMS. Bottom, Semitransparent overlays illustrating KORD expression in pDMS for individuals across the A-P axis. A-P coordinates are in millimeters relative to Bregma. **c**, SigAA performance during session 5 of training for rats assigned to receive Vehicle versus SalB in Test 1. Unpaired t-test: t_10_=0.89 p=0.40. **d**, Left, shuttles per WS for Vehicle versus SalB Tests. Two-way ANOVA: F_4,88_=4.36 **p=0.003. Right, shuttles per WS for Block 1 of Tests. Paired t-test: t_11_=2.339 *p=0.039. **e**, Left, latency to first AR for Vehicle versus SalB Tests. Two-way ANOVA: F_4,88_=1.85. Right, latency to first AR during Block 1 of Tests. Paired t-test: t_11_=1.713 p=0.115. **f**, WS freezing for Block 1 of Vehicle versus SalB Tests. Paired t-test: t_11_=2.36 p=0.037. **g**, Top, example of bilateral KORD expression in DLS. Bottom, Semitransparent overlays illustrating KORD expression in DLS for individuals across the A-P axis. A-P coordinates are in millimeters relative to Bregma. **h**, SigAA performance during session 5 of training for rats assigned to receive Vehicle versus SalB in Test 1. Unpaired t-test: t_5_=0.73 p=0.50. **i**, Left, shuttles per WS for Vehicle versus SalB Tests. Two-way ANOVA: F_4,48_=0.11. Right, shuttles per WS for Block 1 of Tests. Paired t-test: t_6_=0.35 p=0.74. **j**, Left, latency to first AR for Vehicle versus SalB Tests. Two-way ANOVA: F_4,48_=0.22. Right, latency to first AR during Block 1 of Tests. Paired t-test: t_6_=0.53 p=0.61. **k**, WS freezing for Block 1 of Vehicle versus SalB Tests. Paired t-test: t_6_=0.66 p=0.53. **l-n**, Diagrammatic summary of results. All data in the figure are shown as means. Error bars (s.e.m.) are shown for between-subject analyses.

### DLS is required for habitual active avoidance

According to dual process accounts of instrumental behavior, goal-directed and habitual associations co-exist after overtraining and are expressed via parallel circuits that include pDMS and DLS^31,32^. Thus, impairing DLS activity may not reduce overt avoidance behavior, as this will simply disinhibit goal-directed responding (Fig. 4l,m). To implicate DLS in habitual responding, a tool like outcome devaluation or contingency degradation is necessary to also undermine goal-directed actions and produce a decrement in overt responding (Fig. 4o).

**Fig. 4.**
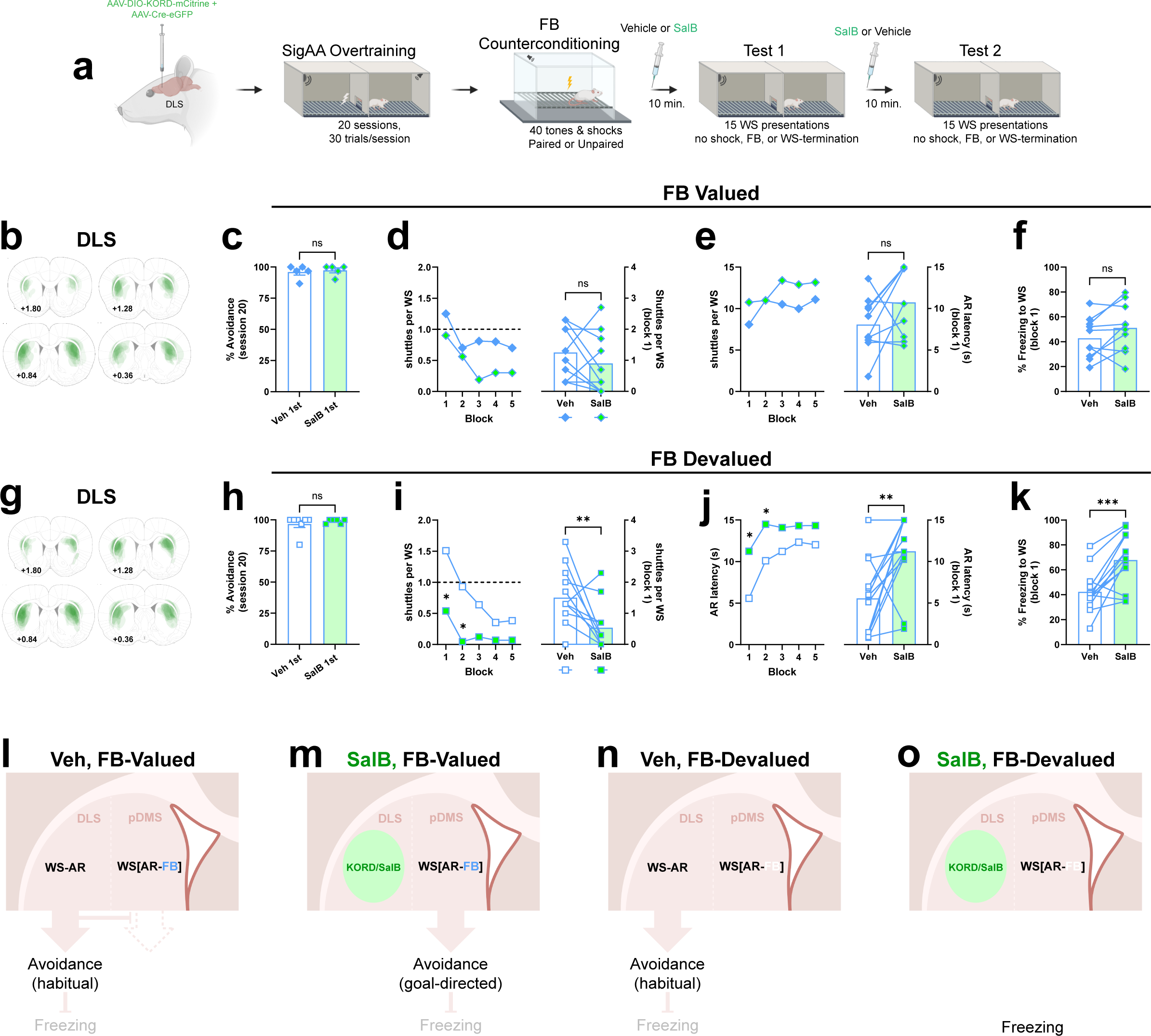
Inhibition of DLS impairs habitual avoidance in male rats. **a**, Schematic of the experimental paradigm. Note that all rats were overtrained and expressed KORD bilaterally in DLS. All rats received 40 tones and shocks during the counterconditioning phase, either unpaired (FB-Valued, n=10) or paired (FB-Devalued, n=13). **b**, Semitransparent overlays illustrating KORD expression in DLS for individuals assigned to FB-Valued condition. A-P coordinates are in millimeters relative to Bregma. **c**, SigAA performance during session 20 of training for FB-Valued rats assigned to receive Vehicle versus SalB in Test 1. Unpaired t-test: t_8_=0.42 p=0.68. **d**, Left, shuttles per WS for Vehicle versus SalB Tests in FB-Valued condition. Two-way ANOVA: F_4,72_=0.57. Right, shuttles per WS for Block 1 of Tests in FB-Valued condition. Paired t-test: t_9_=1.10 p=0.30. **e**, Left, latency to first AR for Vehicle versus SalB Tests for FB-Valued condition. Two-way ANOVA: F_4,72_=0.84. Right, latency to first AR during Block 1 of Tests for FB-Valued condition. Paired t-test: t_9_=1.76 p=0.11. **f**, WS freezing for Block 1 of Vehicle versus SalB Tests in FB-Valued condition. Paired t-test: t_9_=1.83 p=0.10. **g**, Semitransparent overlays illustrating KORD expression in DLS for individuals assigned to FB-Devalued condition. **h**, SigAA performance during session 20 of training for FB-Devalued rats assigned to receive Vehicle versus SalB in Test 1. Unpaired t-test: t_11_=0.71 p=0.49. **i**, Left, shuttles per WS for Vehicle versus SalB Tests in FB-Devalued condition. Two-way ANOVA followed by Šídák’s test: F_4,96_=2.216 (Block x Drug); F_1,24_=21.23 ***p=0.0001 (Drug). Group comparisons, SalB versus Vehicle, Block1 *p=0.03, Block 2 *p=0.02. Right, shuttles per WS for Block 1 of Tests in FB-Devalued condition. Paired t-test: t_12_=3.14 **p=0.009. **j**, Left, latency to first AR for Vehicle versus SalB Tests for FB-Devalued condition. Two-way ANOVA followed by Šídák’s test: F_4,96_=1.43 (Block x Drug); F_1,24_=20.65 ***p=0.0001 (Drug). Group comparisons, SalB versus Vehicle, Block 1 *p=0.02, Block 2 *p=0.03. Right, latency to first AR during Block 1 of Tests for FB-Devalued condition. Paired t-test: t_12_=3.35 **p=0.006. **k**, WS freezing for Block 1 of Vehicle versus SalB Tests in FB-Devalued condition. Paired t-test: t_12_=4.61 ***p=0.0006. **l-o**, Diagrammatic summary of results. All data in the figure are shown as means. Error bars (s.e.m.) are shown for between-subject analyses.

To test whether DLS contributes to avoidance habits, we expressed KORD bilaterally in DLS, overtrained all rats, and used paired counterconditioning to devalue FB tones in half the rats (FB-Devalued group). The remaining rats received unpaired counterconditioning (FB-Valued group). As above, rats then received two Tests consisting of 15 WS presentations after counterbalanced Vehicle or SalB injections. At the end of SigAA overtraining, rats assigned to receive Vehicle- or SalB-first displayed equally high levels of avoidance responding (Fig. 4c,h). In the FB-Valued group, SalB had no effect on WS-shuttles, AR latencies, or WS-freezing (Fig. 4d-f). However, in the FB-Devalued group, SalB impaired WS-shuttles, increased AR latencies, and increased WS-freezing (Fig. 4i-k). This pattern of results supports the hypothesis that parallel circuits in dorsal striatum control goal-directed vs. habitual ARs, with DLS being critical for habitual ARs (Fig. 4l-o).

### FB-Devaluation fails to impair avoidance in female rats

Male data are reported in Fig. 2 and Extended Fig. 2, showing a sensitivity to FB devaluation after moderate training, but not overtraining. An identical analysis in females shows insensitivity to FB devaluation, regardless of the amount of training. Females assigned to FB-Devalued or FB-Valued groups showed equally high avoidance rates at the end of SigAA training (Fig. 5b,h). During counterconditioning, tone-freezing rose quickly in FB-Devalued rats but remained low in FB-Valued rats (Fig. 5c,i), similar to male rats (Fig. 2c,i). During the Devaluation tests, WS-shuttles (Fig. 5e,k), AR latencies (Fig. 5f,l), and WS-freezing (Fig. 5g,m) were similar in both groups.

**Fig. 5.**
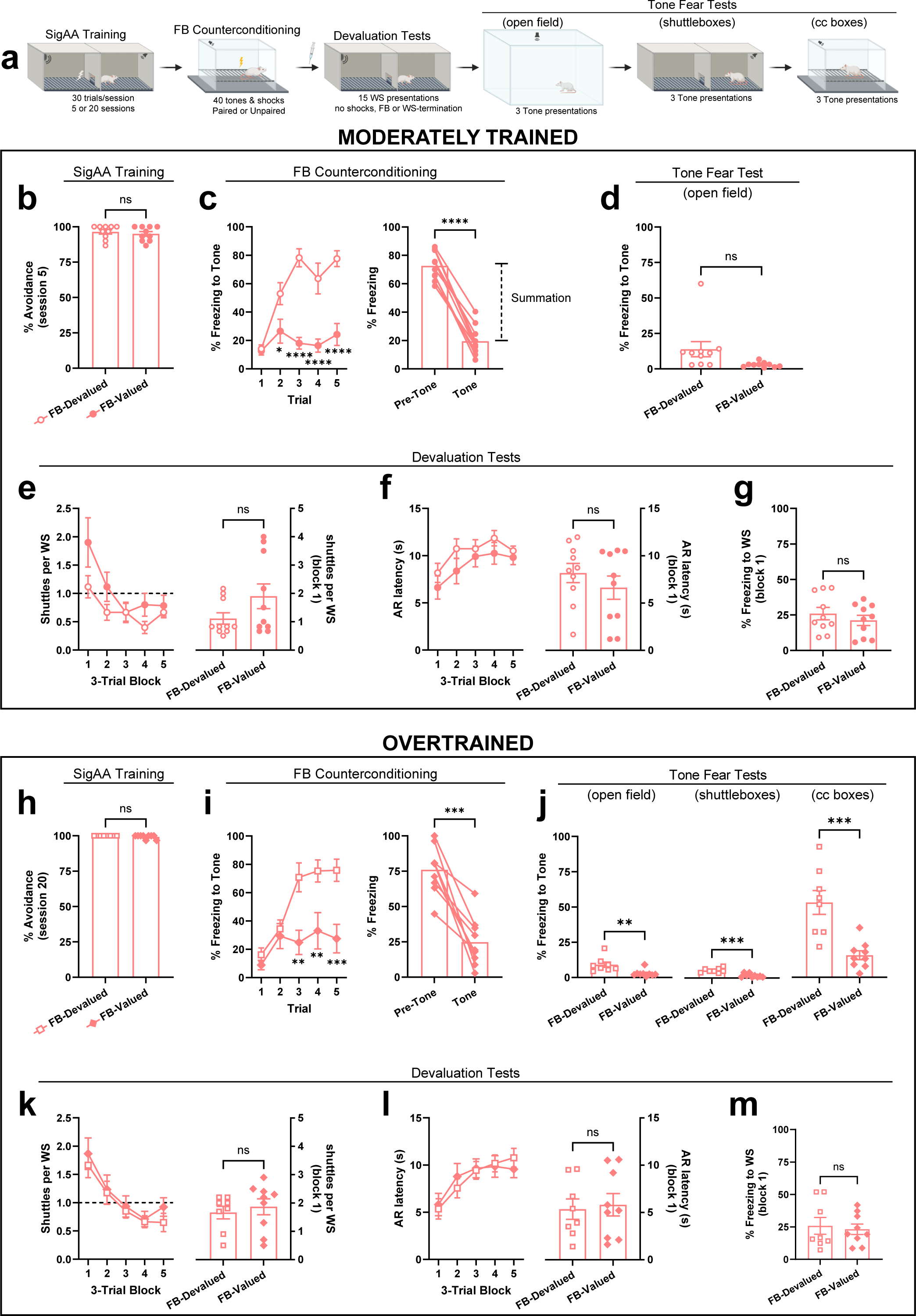
Devaluation of response-produced safety signals after moderate training versus overtraining in female rats. **a**, Schematic of the devaluation paradigm. Panels **b-g**: moderately trained rats. Panels **h-m**: overtrained rats. Note that moderately trained rats received only one Tone Fear Test in the open field. Additional Tone Fear Tests in the counterconditioning boxes and shuttleboxes were later added for overtrained rats to address differences between males and females. **b**, SigAA performance during session 5 of training for rats assigned to FB-Devalued (n=10) and FB-Valued (n=10) groups. Unpaired t-test: t_18_=0.61 p=0.55. **c**, Left, tone freezing during the first 5 trials of counterconditioning. Two-way ANOVA followed by Šídák’s test: F_4,72_=9.183. Group comparisons, FB-Valued versus FB-Devalued, Trial 2 *p=0.03, Trial 3 ****p<0.0001, Trial 4 ****p<0.0001, Trial 5 ****p<0.0001. Right, Pre-Tone versus Tone freezing during first 5 trials of counterconditioning for FB-Valued rats. Paired t-test: t_9_=17.11 ****p<0.0001. **d**, Tone freezing in the novel open field. Unpaired t-test: t_18_=2.06 p=0.054. **e**, Left, shuttles per WS for both Devaluation Tests. Two-way ANOVA: F_4,72_=1.77. Right, shuttles per WS for Block 1 of Devaluation Tests. Unpaired t-test: t_18_=1.64 p=0.12. **f**, Left, latency to first AR for both Devaluation Tests. Two-way ANOVA: F_4,72_=0.38. Right, latency to first AR during Block 1 of Devaluation Tests. Unpaired t-test: t_18_=0.97, p=0.345. **g**, WS freezing for Block 1 of Devaluation Tests. Unpaired t-test: t_18_=0.85, p=0.41. **h**, SigAA performance during session 20 of overtraining for rats assigned to FB-Devalued (n=8) and FB-Valued (n=9) groups. Unpaired t-test: t_15_=1.42 p=0.18. **i**, Left, Tone freezing during the first 5 trials of counterconditioning. Two-way ANOVA followed by Šídák’s test: F_4,60_=3.85 **p=0.008. Group comparisons, FB-Valued versus FB-Devalued, Trial 3 **p=0.002, Trial 4 **p=0.005, Trial 5 ***p=0.0009. Right, Pre-Tone versus Tone freezing during first 5 trials of counterconditioning for FB-Valued rats. Paired t-test: t_8_=6.39 ***p=0.0002. **j**, Tone freezing in the novel open field (left), shuttleboxes (middle), and counterconditioning boxes (right). Unpaired t-tests: t_15_=3.20 **p=0.006 (left); t_15_=5.21 ***p=0.0001 (middle); t_15_=4.33 ***p=0.0006 (right). **k**, Left, shuttles per WS for both Devaluation Tests. Two-way ANOVA: F_4,60_=0.33. Right, shuttles per WS for Block 1 of Devaluation Tests. Unpaired t-test: t_15_=0.56 p=0.58. **l**, Left, latency to first AR for both Devaluation Tests. Two-way ANOVA: F_4,60_=0.81. Right, latency to first AR during Block 1 of Devaluation Tests. Unpaired t-test: t_15_=0.29 p=0.78. **m**, WS freezing for Block 1 of Devaluation Tests. Unpaired t-test: t_15_=0.36 p=0.73. Data in **c** (right) and **i** (right) are shown as mean + individuals. All other data in the figure are shown as mean ± s.e.m.

### Counterconditioning is highly context-dependent in female rats

Note that tone-freezing data during counterconditioning shows that our FB devaluation protocol was effective in both sexes (compare Fig. 5c,i to Fig. 2c,i). Thus, there are at least three possible explanations for the failure to impair moderately trained avoidance with FB devaluation in females: 1) females transition to habits sooner, 2) females rely on a different outcome for goal-directed avoidance (e.g., US omission), or 3) counterconditioning of safety signals does not transfer to other contexts in females. Tone Fear Tests strongly support the third explanation. Moderately trained females subjected to FB devaluation showed very low tone-freezing in the open field (14%; Fig. 5d). To explore this further, we added additional Tone Fear Tests in the shuttleboxes and the counterconditioning boxes for overtrained rats. Although FB-Devalued females showed more tone-freezing than FB-Valued females in all locations (Fig. 5j), average freezing was extremely low in the open field (FB-Devalued: 9%, FB-Valued: 3%) and shuttleboxes (FB-Devalued: 5%, FB-Valued: 2%). The only place where females showed substantial tone-freezing after FB devaluation was in the counterconditioning boxes (53%; Fig. 5j, right). Thus, unlike males (Fig. 2d,j), females only express counterconditioning of safety signals in the context where it occurred.

To circumvent the context-dependence of counterconditioning in females, we ran three additional experiments where females were moderately trained, and counterconditioning occurred in the shuttleboxes. In the first experiment, counterconditioning was identical to previous experiments except it occurred in the shuttleboxes (Extended Fig. 5a-f). This appeared to solve the context-dependence problem, as FB-Devalued rats showed high tone-freezing in the shuttleboxes but not the open field. However, avoidance collapsed during the Devaluation Tests and there were no group differences. The second experiment duplicated the first except we used a weaker counterconditioning procedure (20 tones and shocks, paired or unpaired) to maintain avoidance responding for FB-Valued rats (Extended Fig. 5g-l). This produced nearly identical results. In the third experiment, we again used the weaker counterconditioning procedure, but closed the shuttlebox door during counterconditioning (Extended Fig. 5m-r). This was done because we noticed some rats attempting to shuttle during the counterconditioned tone, and it is unclear how this affects WS-shuttling at test. Although this led to more ARs during the Devaluation Tests, we again observed no group differences. Interestingly, FB-Devalued females showed high tone-freezing in the shuttleboxes, but only when the door was closed – as during counterconditioning (Extended Fig. 5r). Together, these data indicate a remarkable context-dependence of safety signal counterconditioning in female rats.

### Degrading AR-safety contingency impairs moderately trained avoidance in both sexes

In appetitive studies, goal-directed responding is also sensitive to A-O contingency degradation (CD). For instance, rats barpressing for food will reduce responding if food is delivered for free (non-contingently), even if food remains available upon pressing^33^. We designed a CD protocol using male rats shuttling in an unsignaled active avoidance (USAA) paradigm^26^. During USAA, there is no explicit WS and shocks are scheduled for delivery every 5s. Each shuttle produces a FB cue and delays footshock by 30s. Shuttles during response-shock intervals are coded as ARs and shuttles following a shock are coded as escape responses (ERs). Explicit response-produced FB cues improve learning^34^ and become safety signals with USAA training^35^. After reaching training criterion, subjects were matched into equal Control (CTL) and CD groups. CTL rats received four additional USAA training sessions. CD rats received similar sessions, except additional safety signals (+30s of safety from shock) were delivered non-contingently (Fig. 6a). This reduced ARs and ERs by about 50% (Fig. 6b,c) while having no significant effect on the frequency of shocks (Fig. 6d). By design, CD rats also received significantly more safety signals than CTL rats (Fig. 6e). These data converge with our FB devaluation data (Fig. 2), further implicating safety signals in goal-directed avoidance.

**Fig. 6.**
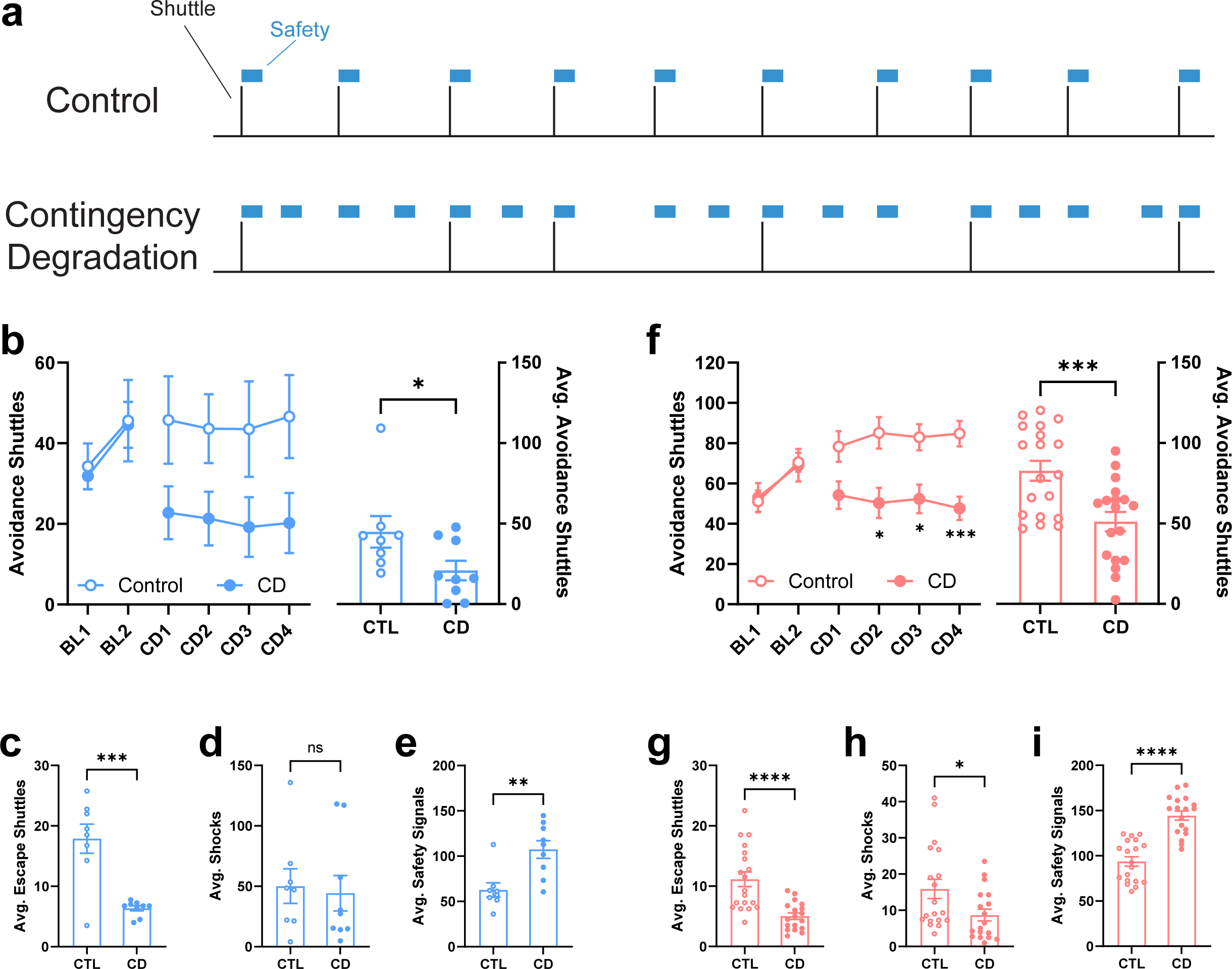
Contingency degradation impairs goal-directed avoidance in male and female rats. **a**, Schematic of the contingency degradation protocol. **b**, Left, Number of avoidance shuttles during the last two training sessions (BL1-2) and four contingency degradation sessions (CD1-4) for male rats tested at TAMU. Rats assigned to Control (CTL; n=8) and CD (n=9) groups performed similarly during the baseline phase. Two-way ANOVA: F_1,15_=0.02 p=0.89. CD rats avoided less than CTL rats during the contingency degradation phase. Two-way ANOVA: F_3,45_=0.09 (Session x Group), F_1,15_=4.57 *p=0.049. Right, avoidance shuttles per CD session. Unpaired t-test: t_15_=2.14 *p=0.049. **c**, Average escape shuttles per CD session. Unpaired t-test: t_15_=4.98 ***p=0.0002. **d**, Average shocks delivered per CD session. Unpaired t-test: t_15_=0.28 p=0.78. **e**, Average safety signals per session. Unpaired t-test: t_15_=3.52 **p=0.003. **f**, Left, Number of avoidance shuttles during the last two training sessions (BL1-2) and four contingency degradation sessions (CD1-4) for female rats tested at TAMU and NKI (combined here, see Extended Fig.6 for separate results). Rats assigned to Control (n=19) and CD (n=18) groups performed similarly during the baseline phase. Two-way ANOVA: F_1,35_=0.15. CD rats avoided less than Control rats during the contingency degradation phase. Two-way ANOVA followed by Šídák’s test: F_3,105_=1.07 (Session x Group), F_1,35_=13.34 ***p=0.0008 (Group). Group comparisons, CD versus CTL, CD3 *p=0.01, CD4 *p=0.01, CD5 ***p=0.0005. Right,Average avoidance shuttles per CD session. Unpaired t-test: t_35_=3.65 ***p=0.0008. **g**, Average escape shuttles per CD session. Unpaired t-test: t_35_=4.54 ****p<0.0001. **h**, Average shocks per CD session. Unpaired t-test: t_35_=2.28 *p=0.03. **i**, Average safety signals per CD session. Unpaired t-test: t_35_=7.01 ****p<0.0001. All data in the figure are shown as mean ± s.e.m.

CD may be ideal for assessing goal-directedness of ARs in females because it preserves the value of the outcome (FB=safety) and occurs in the training context. To test whether females are also sensitive to CD after moderate USAA training, parallel experiments were conducted at two institutes by different experimenters using identical protocols and equipment. Results are combined in Fig. 6f-g and shown separately in Extended Fig. 6. Like males, CD females received more safety signals and emitted significantly fewer ARs and ERs (Fig. 6i,f,g). CD females also received fewer shocks than males (Fig. 6h), likely because they shuttle at a higher rate throughout the sessions. Remarkably similar patterns of ARs, ERs, shocks, and safety signals were observed at both institutes (Extended Fig. 6), indicating that this CD effect is robust. These data suggest that females also rely on FB safety signals for goal-directed avoidance after moderate training.

## Discussion

Our results indicate that response-produced FB stimuli are transformed into safety signals during AA training that support goal-directed avoidance. They also show that overtrained ARs are insensitive to FB devaluation. These data confirm that AA can indeed produce instrumental responses directed towards safety, a topic that has been long debated^4,11,36,37^ but, until now, never demonstrated empirically. These findings, along with our KORD/SalB studies, are also consistent with associative dual process accounts of instrumental behavior implicating parallel dorsal striatum circuits in goal-directed vs. habitual responding^9,31,38,39^. This pattern of results suggests that AA relies on similar psychological and neural mechanisms as other positively reinforced instrumental behaviors once FB stimuli become valued safety signals. Our data also confirm that freezing and ARs are anticorrelated, suggesting that AA is an anxiety-related response to threat that is distinct from fear responses and selected in contexts where safety is attainable. These results challenge traditional theories that frame avoidance behavior as reflexive (S-R), negatively reinforced, and fear motivated.

A core finding in appetitive instrumental research is that positively reinforced behavior controlled by goal-directed vs. habitual associations depends on parallel, competitive circuits in dorsal striatum. pDMS is required for devaluation-sensitive responses and DLS is required for devaluation-insensitive responses^17,19,31,40^. Little is known about the circuits mediating AA. However, several reports are consistent with the idea that AA is reinforced by safety and controlled by different brain circuits after overtraining. AA learning and performance also depends on basolateral amygdala^26,41–43^, and on the serial connection between basolateral amygdala and nucleus accumbens shell (NAcS)^25^. However, overtrained ARs are amygdala-independent^26,42^. Avoidance behavior can also be bidirectionally modulated by manipulating dopamine prediction error signals to AR-FB in nucleus accumbens after moderate training, but not overtraining^44–46^. NAcS has also been implicated in the positive reinforcement of overtrained ARs by safety^47^. Finally, Darvas and colleagues found that replacing dopamine in amygdala+striatum rescued the ability of dopamine-deficient mice to acquire ARs^48^. However, once learned, dopamine was required only in striatum to perform the response.

Studies of pDMS in goal-directed avoidance are sparse and inconclusive. Operating on the assumption that goal-directed associations are acquired more rapidly than habitual associations^18^, we tested the hypothesis that pDMS is required for avoidance after moderate training – a phase we found to be sensitive to FB-devaluation. KORD-mediated inhibition of pDMS reduced ARs during the WS and caused a return of WS-induced freezing after moderate training. Interestingly, AR performance was unimpaired when aDMS or DLS were suppressed after moderate training, or when pDMS was suppressed after overtraining. This anatomically-specific and training-sensitive effect provides strong support that goal-directed avoidance depends on the same posterior region of DMS that mediates appetitively reinforced instrumental actions^17^. Also consistent with appetitive studies^19,32^, inhibition of DLS impaired avoidance only in rats that also received FB-Devaluation. This suggests that avoidance relies on the same competitive circuits in dorsal striatum that mediate instrumental actions vs. habits established with rewards. These findings also demonstrate the utility of our FB devaluation paradigm for identifying goal-directed ARs and isolating avoidance habits for study.

Although negative reinforcement theories dominated during the heyday of AA research, the idea that safety signals may positively reinforce ARs was introduced as early as 1936 by Konorski and Miller. This was echoed by Soltysik, who called inhibitory AR-FB a “reward” that “stabilizes avoidance behavior and makes it virtually independent of the noxious US”^12^. Studies since then support the notion that safety signals reinforce ARs without directly demonstrating an AR-FB association^10,15,34,49–51^. Initial work attempting to distinguish between avoidance actions vs. habits focused on devaluing aversive USs^3,52–55^. However, none show that AR decrements occur from the beginning of devaluations tests, when responding is most clearly linked to a memory of the outcome value^24^. Also, the amount of training was typically not varied, and no studies show a loss of devaluation effects with overtraining. Although these data suggest that current US value contributes to avoidance, it is possible that devaluing an aversive US affects ARs indirectly by reducing the value of safety cues that predict its absence. Indeed, some have described the conditioned inhibition that defines safety signals as a “slave process” to aversive excitation^56^.

We directly tested the safety signal hypothesis of goal-directed avoidance by creating a novel outcome-devaluation procedure. The near complete collapse of avoidance after devaluation suggests that response-produced safety signals are the primary outcome supporting AR performance after moderate training. These results may help to resolve problems that slowed progress in understanding AA. The finding that safety signals reinforce avoidance may explain how WS-termination and US-omission contribute to avoidance behavior^11^. These events are inversely related to AR-FB cues and likely both support Pavlovian safety learning^[2]^. They also help explain the extreme resistance to extinction sometimes observed after avoidance training^57^. Discontinuing the US does nothing to undermine AR◊FB or WS◊AR associations that should maintain responding. Our results are also consistent with many studies observing that measures of fear are poorly correlated with avoidance^4,6^. As ARs are successfully acquired, the meaning of the WS likely changes from a threat of imminent harm to a threat of harm in a context where safety is easily attainable^4^. This should induce a shift from a central state of fear to one of anxiety, where harm is less likely and volitional action is possible^58,59^. Since fear and anxiety are distinct defensive brain states^60^, innate fear reactions like freezing are suppressed or remain untriggered during anxiety states that support avoidance. However, if an effective AR is blocked or otherwise impaired, the fear state returns along with inflexible reactions like freezing^61^ (see also Figs. 2,3,4).

Although avoidance behavior was very similar in males and females, we found a major sex difference in our FB-Devaluation studies. Females showed no AR decrement when the FB-tone was counterconditioned after moderate training. This was not due to a failure of safety conditioning or a failure of counterconditioning. Instead, females displayed a remarkable degree of context dependence in their expression of counterconditioning that was not observed in males. We cannot find any previous reports of this sex difference in the literature. However, studies on a different form of counterconditioning were conducted primarily in females and the expression of counterconditioning was context-dependent^62^. As a practical matter, our result suggests that FB-Devaluation may not be a useful tool for distinguishing goal-directed and habitual ARs in female rats.

In our final experiment, we used a novel contingency degradation protocol after USAA training to confirm our behavioral findings in males and determine if females also rely on safety signals for goal-directed avoidance. After moderate training, males adjusted their AR rates downward when free (non-contingent) FB safety signals were presented during training. This was replicated in females by different experimenters at different institutions, suggesting the effect is robust. Thus, contingency degradation may be especially useful for studying neural circuits of goal-directed vs. habitual avoidance in females. However, more work is needed to determine if delivering free safety signals loses effectiveness with overtraining.

In summary, this work suggests that volitional action is indeed possible under threat, even intense threat^63^, provided that responses are available in the context to attain safety^58^. Our safety-signal devaluation protocol provides a new tool for differentiating between goal-directed and habitual ARs in neurobiological studies and in studies of aberrant avoidance habits in humans. Recent work suggests that patients with OCD^3^, or with a history of early life stress^54^, have slightly stronger avoidance habits. It would be interesting to see if evidence for more robust AR habits is obtained after safety signal devaluation. Our data showing strong context-dependence of counterconditioning in females may also have clinical implications if replicated in humans. Experiences that devalue an avoidance reinforcer may only reduce avoidance in the contexts where they happen, possibly contributing to the higher rate of anxiety disorders characterized by avoidance in women. Interestingly, a recent study found that human ratings of “relief pleasantness” during response-produced safety periods relate to reward prediction error signals that decline as avoidance is acquired^64^. However, those scoring low on distress tolerance showed no such decline, suggesting possible over-reinforcement of avoidance habits. A safety signal devaluation test may help resolve this question and potentially identify a new mechanism contributing to anxiety. Lastly, our results provide empirical support for reinforcement learning models that rely on response-produced safety signals to explain AA^65^.

## Methods

### Rats

All procedures were performed in accordance with the National Institute of Health Guide for the Care and Use of Experimental Animals and were approved by the NKI and TAMU Animal Care and Use Committees. Most studies were conducted at NKI with a subset conducted at TAMU (Fig. 6 & Extended Fig. 6). Subjects were female and male Sprague-Dawley adult rats weighing 250-300g upon arrival (NKI: Hilltop Laboratory Animals, Scottsdale, PA, USA; TAMU: Envigo, Indianapolis, IN, USA). Rats were pair-housed in transparent plastic high-efficiency particulate absorption (HEPA)-filtered cages with cob bedding on a 12:12h (NKI) or (14:10) light-dark cycle within temperature and humidity-controlled environments. Food and water were available ad libitum throughout the duration of the experiment. Animals were allowed a minimum of five days to acclimate to the vivarium before all experiments. Following surgery, animals were singly housed for a minimum of one week and returned to pair-housing once recovered.

### Shuttlebox apparatus

All equipment was from Coulbourn Instruments. SigAA training and devaluation tests occurred in two-way shuttle boxes (Model H10-11R-SC) within sound-attenuating chambers (SAC Model: H10-24A). Shuttle chambers were divided into two identical sections (25.4 depth x 50.8 width x 30.5 cm height) by metal dividers which contained a small passage (8.0 cm wide, 8.5 cm high) that allowed rats to move freely between sides. In one experiment, the passage connecting the shuttlebox chambers was blocked (door closed) during counterconditioning and one Tone Fear Test via a metal insert held in place with magnets. A houselight with small 0.5 W bulbs in each compartment dimly illuminated the shuttle-boxes (25 cm high). A PC computer installed with Graphic State software (v. 3 or 4) and connected to the chambers via the Habitest Linc System (Model H01-01) delivered 80dB white noise warning stimuli (WSs), feedback (FB) tones (5kHz, 80dB) and shock unconditioned stimuli (USs) during behavioral sessions. WS and FB were produced by an audio generator (Model A12-33) and presented through speakers (Model H12-01R) located on opposing walls in each chamber (20 cm high). Scrambled shock (Model H13-15) was delivered to a grid floor consisting of 32 stainless steel bars arranged in parallel to the dividers (Model H10-11R-XX-SF). Shuttling was automatically registered by infrared beams and recorded throughout the sessions (Model H20-95-X). Behavior was monitored by security video cameras mounted on the ceiling of each shuttlebox chamber and recorded to DVD.

### Counterconditioning apparatus

All equipment was from Coulbourn Instruments. Counterconditioning and some Tone Fear Tests took place in rodent conditioning chambers (28.5 depth x 26 width x 28.5 height cm, Model H10-11R-TC) enclosed by sound-attenuating cubicles (Model H10-24A, Coulbourn Instruments) and dimly lit by a houselight with a 0.5 W bulb (25 cm high). A computer, installed with Graphic State software (v. 3) and connected to the chambers via the Habitest Linc System (Model H01-01) delivered tones (5s, 5kHz, 80dB) and shock US stimuli during behavioral sessions. Tones were produced by an audio generator (Model A12-33) and presented through a single speaker (Model H12-01R) located in the front panel of the chamber (20 cm high). Scrambled shocks (Model H13-15) were delivered to a grid floor containing 18 stainless steel bars arranged in parallel (H10-11R-TC-SF). Animal movement was monitored by a security video camera mounted on the ceiling of the cubicle and recorded to DVD.

### 2-way SigAA training procedure

Each session began in illuminated shuttleboxes with a 5-minute stimulus-free acclimation. On the first day of training only, after acclimation, rats received a single Pavlovian (WS-US paired) trial with an inescapable 0.5s footshock US, followed by 30 SigAA trials. The remaining days of training (day 2+) did not include the initial Pavlovian trial. After the acclimation period on subsequent training days (day 2+), rats received 30 SigAA trials with an average intertrial interval (ITI) of 2 minutes (random assignment without replacement, range 90-150s). Trials consisted of an 80dB white noise WS presentation that lasted a maximum of 15 s. Shuttling during the WS led to immediate WS termination, prevented delivery of the scheduled footshock US (0.5 s; 1.0 mA males, 0.7 mA females), and triggered the delivery of a pure tone FB cue (5s, 5kHz, 80dB), or in the case of FB-controls (see below), a 5s period of darkness (houselights turned off). If no avoidance shuttle was performed during the 15s WS, the inescapable US was delivered for 0.5s and no FB was delivered. Avoidance responses (ARs) were defined as shuttles during the WS. After 30 avoidance trials were completed, animals were given a 60s final period in the shuttle box before removal. Training (one session/day) continued for either 5 days (moderately trained) or 20 days (overtrained) with 5 sessions occurring per week. Rats failing to reach 70% successful avoidance by session 5 (poor avoiders) were removed from experiments.

### Counterconditioning procedure

During the last five days of avoidance training (days 1-5 for moderately-trained rats, days 16-20 for overtrained rats), animals were placed in the illuminated counterconditioning apparatus and given 30 min. stimulus-free habituation sessions. It is our impression that this habituation phase is critical for obtaining effective counterconditioning of safety signals. 1d after the final SigAA training session, animals were placed in the illuminated counterconditioning boxes for a 3 min. acclimation period. FB-Paired and FB-control rats received 40 trials of the 5s tone (5kHz, 80dB) that co-terminated with a 0.5s footshock US (0.7mA for females, 1.0mA for males; avg. ITI: 180s, random assignment without replacement, range 150-210 s). For the FB-control group, the tone was novel since they had been SigAA-trained with a visual FB cue. Unpaired rats received 40 tones and shocks in an explicitly unpaired fashion. Trial 1 began with a footshock, with tones and shocks alternating thereafter. Shocks occurred on average every 3 minutes, with placement closer to the onset of the next tone than the termination of the previous tone (to prevent trace conditioning). Shock-to-tone intervals ranged from 28-88s (mean: 58s) and tone-to-shock intervals ranged from 87-147s (mean: 117s). All intervals were selected from lists at random without replacement. Total session times for all animals were approximately 125 min. Two experiments with female rats conducted counterconditioning in the shuttleboxes using only 20 tones & shocks (Paired versus Unpaired, Extended Fig. 5). These sessions were the same as the standard procedure except the total session duration was shorter (∼65 min).

### Devaluation/AR Tests

Note that all Devaluation or AR Tests were conducted 10 min. after i.v. injections with 1 ml/kg injection volumes. For within-subject KORD studies, rats were injected with Veh or SalB (counterbalanced). For other purely behavioral experiments, rats were injected with Veh before both tests. This was done to control for stress before testing in all studies, and to allow team members to practice i.v. injections before the critical KORD studies. After acclimation in the shuttleboxes (SigAA training context) for 5 min., rats received 15 presentations of the 80dB, 15s WS (mean ITI: 120s, random assignment without replacement, range 90-150s). Shuttling was monitored throughout the session but had no effect on WS duration (no WS-termination). There were also no FB presentations (tones or houselight-off) or footshock US presentations. Devaluation/AR tests occurred twice, with 1d off between tests.

### Tone fear tests

After all devaluation tests were completed, rats from behavioral experiments were placed in a novel open field (90 x 90 x 40 cm) for a 2 minute acclimation phase followed by 3 presentations of a 5 kHz, 80 dB, 60s tone cue (1 min, it is; Figs. 2d, 2j, 5d, 5j, and Extended Figs. 5f,l,r). Video was recorded to DVD using a digital camera and freezing responses were rated offline. Some rats were also tested in the two-way shuttleboxes where the avoidance acquisition training originally occurred using an identical protocol with the shuttlebox door open or closed (Figs. 2j, 5j, and Extended Fig. 5f,l,r). Some rats were subsequently tested in the counterconditioning boxes (Figs. 2j, 5j) using an identical protocol. Tone fear tests occurred 2-3d after the final Devaluation Test, with only one test/day (2-3d interval between tests).

### USAA and Contingency Degradation procedures

Rats were placed in the shuttlebox, with offset of the houselight indicating the beginning of the session. Shock (1.0 mA for males, 0.7 mA for females) was delivered every 5s (shock-shock, or S-S interval) unless the subject shuttled to the opposite chamber. Every shuttle earned a FB tone (0.5 s, 70 dB, 2kHz) and a 30s shock-free period (response-shock, or R-S interval). Shuttles during S-S intervals were considered escape responses (ERs) and shuttles during R-S intervals were considered avoidance responses (ARs). If the subject failed to shuttle during S-S or R-S intervals, shocks resumed every 5 s. All subjects received USAA until reaching “good avoider criterion” (2 consecutive sessions with 20 or more ARs)^26^. Rats failing to reach this criterion by session 10 were deemed poor avoiders and eliminated from the study. Session duration was 25 minutes.

Contingency Degradation (CD) phase: Following USAA training, animals were nonrandomly assigned to Control or CD groups based on comparable levels of criterion performance. Control animals received four more days of USAA training, while experimental animals received four days of USAA-CD. For the latter, all parameters were the same as the USAA protocol except during the shock-free period (regardless of whether it was earned by escape or avoidance), every 5s there was a 1/3 chance that the rat would receive a ‘free’ SS (and additional shock-free period) in a way that was not contingent on behavior.

### Surgery for DREADD experiments

Rats were anesthetized with a mixture of ketamine (100 mg/kg, i.p.) and xylazine (10 mg/kg, i.p.) and placed in a stereotaxic apparatus (David Kopf Instruments). Supplemental doses of the mixture were given as needed to maintain a deep level of anesthesia. Brain areas were targeted using coordinates from the Paxinos and Watson Brain Atlas. The skull was exposed and small burr holes were drilled above the target area. Rats were bilaterally injected with 0.5 µL AAV using a 1.0 µL Hamilton Neuros syringe (Hamilton Co.) at a rate of 0.1 µL/min. Following infusion, the syringe was left in place for an additional 5-8 minutes to allow fluid diffusion away from the needle tip before withdrawing. AAV injections consisted of a 1:10 cocktail of Cre [AAV5-CMV-HI-eGFP-Cre.WPRE.SV40 (Addgene plasmid no. 105545) packaged at UPenn Vector Core 2.5 × 1014 GC ml−1] and KORD [AAV8-HSyn-DIO-HA-KORD-IRES-mCitrine (Addgene plasmid no. 65417), packaged at UPenn Vector Core, 8.9 × 1013GC ml−1], respectively. Injections were made into the anterior dorsomedial striatum (aDMS; y = +0.5 mm anterior to Bregma, x = +/- 2.1 mm lateral to midline, z = - 4.5 mm ventral from skull surface), the posterior dorsomedial striatum (pDMS; y = -0.12 mm, x = +/- 2.45 mm, z = -4.9 mm), or the dorsolateral striatum (DLS; y = +0.75 mm, x = +/- 3.5 mm, z = -5.1, -5.6 mm). Immediately following surgery, animals were sutured (Ethicon Vicryl, Fisher Scientific), injected with analgesic (Buprenorphine-ER (s.c., 0.6-1.2 mg/kg), Wedgewood Pharmacy) and allowed to recover on a heat pad. Once awake and alert, rats were returned to the vivarium in a fresh, clean cage to recover for one week singly housed. After one week, rats were returned to pair housing for at least 14 additional days before behavioral training.

### Drug injections

Salvinorin B (SalB, 2.5 mg/mL, Sigma) was dissolved in Vehicle (7% DMSO and 50% PEG in ddH2O) under low heat. Animals were restrained briefly and intravenously injected (i.v., lateral tail vein) with Vehicle or 2.5 mg/kg SalB (1ml/kg volume) 10 min prior to all devaluation/AR Tests. Treatments (Vehicle versus SalB) and order of treatment (Vehicle first versus SalB first) were counterbalanced for all KORD studies evaluating DMS and DLS. Animals with poor or unsuccessful injections were removed from the study.

### Implant surgery for electrophysiology

3-4 weeks after unilateral DMS viral transfection using techniques and coordinates described above, 4 rats were surgically anesthetized with isoflurane, and a topical local anesthetic (1% xylocaine) was administered locally around the wound site. Temperature was maintained at 37°C throughout the surgery using a heating pad. Animals were placed in a stereotaxic device and the skull was exposed. A 2-channel telemetry transmitter (HD-X02, Data Sciences International) was inserted under the skin of the back and stainless-steel electrodes (100µ diameter, Teflon coated, A-M Systems) connected to the telemetry leads were implanted in the DMS bilaterally targeting the region of KORD transfection in the infected hemisphere and the same location in the contralateral uninfected hemisphere. Reference electrodes were implanted in the posterior cortex. Electrodes were attached to the skull with bone-anchor screws and dental cement and the scalp sutured closed. Following recovery from anesthesia, animals were singly housed and allowed to recover for at least 1 week before recording.

### *In vivo* recording and analysis

Recordings were performed in the animal’s home cage by placing the cage on a telemetry receiver. Bilateral local field potentials were recorded in the DMS (amplification 200x, low-pass filtered at 200Hz) and digitized (1000Hz) for recording and analysis with Spike2 software (CED, Inc). Following at least 15 min of baseline recording, rats were injected with SalB (2.5 mg/kg, 1ml/kg, i.v.) and returned to their home cage (<5 min post-injection) for continued recording of spontaneous activity for at least 30min. LFP data were analyzed with FFT (2.4Hz/bin) from at least 10 min. of continuous data pre-injection and starting at least 15 min. post-injection. The post-injection FFT spectrum was normalized to the baseline recording from that same hemisphere, providing a measure of SalB induced change in the infected DMS and the control, uninfected DMS within animals. Following the recording, animals were perfused for histological confirmation of viral expression and electrode placement. In one animal the electrode in the control hemisphere missed the DMS and entered the lateral ventricle, thus only data from the infected hemisphere of this animal was used in the analysis (Extended Fig. 3a-d).

### Histology and immunohistochemistry

After behavioral testing was completed, DMS and DLS animals were anaesthetized with an overdose mixture of ketamine (100 mg/kg, i.p.) and xylazine (10 mg/kg, i.p.) and perfused transcardially with 4% paraformaldehyde (PFA) in 0.01 M phosphate-buffered saline (PBS). The brains were fixed in 4% PFA solution for 24 hours, then post-fixed in 30% sucrose at 4°C for 48-72 hours. Brains were then blocked and sectioned on a microtome (SM2000R, Leica) at 40 µm. These sections were kept in 0.01 M PBS + 0.05% sodium azide (NaAz) until immunohistochemistry. Brain sections were blocked in 1% bovine serum albumin (BSA; Sigma) in 0.1 M PBS for 30-60 minutes at room temperature (RT) to block non-specific binding. KORD-injected brain sections were incubated overnight at RT in Rabbit Anti-GFP Polyclonal Primary Antibody (Invitrogen, #A-11122) in 1% BSA + 0.2% Triton X-100 in 0.01 M PBS (1:1,000) for verification of viral expression in neurons. Following primary antibody incubation, sections were rinsed with agitation three times for 5 minutes each in 0.01 M PBS at RT. Sections were then incubated in Goat Anti-Rabbit IgG (H+L) Cross-Adsorbed Secondary Antibody, Alexa Fluor 488 (Invitrogen, #A-11008) in 0.01 M PBS (1:200) at RT for 30-60 minutes. In some experiments primary and secondary antibody steps were combined using Alexa Fluor 488-tagged GFP Polyclonal Antibody, (Invitrogen, #A21311). After secondary incubation, sections were rinsed with agitation three times for 5 minutes each in 0.01 M PBS at RT, mounted on gelatin-coated slides, cover-slipped in Permount (Fisher, #SP-15) or Prolong Gold Antifade Reagent (Invitrogen, #P36930) and allowed to cure overnight at RT. Slides were placed at 4°C for long-term storage. Sections were imaged using a Leica TCS SP8 confocal microscope (Leica) or Olympus VS120 fluorescent microscope (Olympus). For all experiments, animals were excluded from analysis if virus expression was insufficient, too extensive, or outside regions of interest.

### Assessment of freezing

Videos recorded during counterconditioning and testing sessions were analyzed for freezing by at least one rater who was blind to treatment condition when possible. Blinding was not possible for counterconditioning sessions where reactions to shock were obvious. Freezing was defined as cessation of all non-respiratory movement and is presented as a percentage of the interval assessed. For counterconditioning sessions, freezing was scored for the first 5 trials during pre-tone (5s) or tone (Paired: 4.5s, Unpaired: 5s) intervals. For Devaluation and AR Test sessions, freezing was scored during 15s WS presentations for the first 3 trials (Block 1). For Tone Fear Tests, freezing was scored during each 60s tone presentation.

### Statistics and reproducibility

Data are typically presented as mean ± s.e.m, or mean plus individual values. Most data were analyzed using GraphPad Prism (v. 10.0.3) software. Figures were created with GraphPad Prism, Adobe Illustrator, Inkscape 1.2, and Biorender.com. Error bars in graphs represent ± s.e.m for between-subject analyses, error bars are not included for within-subject graphs. Unpaired Students t-tests were used to compare means between two groups. Paired Students t-tests were used for within-subject comparisons at a single timepoint (e.g. Block 1 histograms). For comparing multiple groups at a single timepoint, ordinary One-way ANOVAs were conducted, assuming equal SDs per group, followed by Dunnett’s multiple comparisons test (single pooled variance). For comparing groups across multiple timepoints, Two-way repeated measures ANOVAs were performed with no assumption of sphericity (Geisser-Greenhouse correction). For experiments with more than two groups, this was followed by Dunnett’s multiple comparison test, with individual variances computed for each comparison. For experiments with two groups, this was followed by Šídák’s multiple comparison test, with individual variances computed for each comparison. To determine variance among animals across sessions (Fig. 1d, right panel), eta-squared (η^2^) was calculated to determine effect size, where η^2^ ≤ 0.06, 0.07–0.14, and η2 > 0.14 indicated small, medium, and large effect sizes, respectively. For each animal, a significant large sized session effect (F = 8.37, p < 0.001, η^2^ = 0.17) was apparent. We therefore calculated coefficients of variation (CV) to assess the extent of variability across sessions was significant using a mixed model ANOVA. Here, CV is inversely related to stereotyped behavior. Effect size calculations were completed using R statistical software. P values <0.05 were considered statistically significant.

## Data availability

All the data that support the findings presented in this study are available from the corresponding author upon reasonable request.

## Acknowledgments

The authors thank Andrew Delamater, Vincent Campese, and Cristina Siller-Perez for helpful discussions on the design of the devaluation procedure and interpretation of data. The project described was supported by Award Number R01MH114931 to C.K.C. from the U.S. National Institutes of Health. R.M.S. received support from a Templeton Foundation grant (TWCF0366). The content is solely the responsibility of the author and does not necessarily represent the official views of the National Institute of Health and National Institute of Mental Health.

## Author contributions

RM conducted surgeries and histology, analyzed and interpreted data, and assisted with writing. EA conducted surgeries, histology, perfusions, and data analyses. SS, LL, and DM participated in all aspects of data collection including surgeries, perfusions, histology, behavioral testing, and scoring of freezing. DW conducted the *in vivo* electrophysiology experiment and analyzed data. MA conducted the contingency degradation experiment at TAMU. JM designed the contingency degradation experiment, collected and analyzed behavioral data, and assisted with writing. CC designed the devaluation experiments, collected and analyzed behavioral data, and wrote the paper.

---

**Extended Fig. 1.**
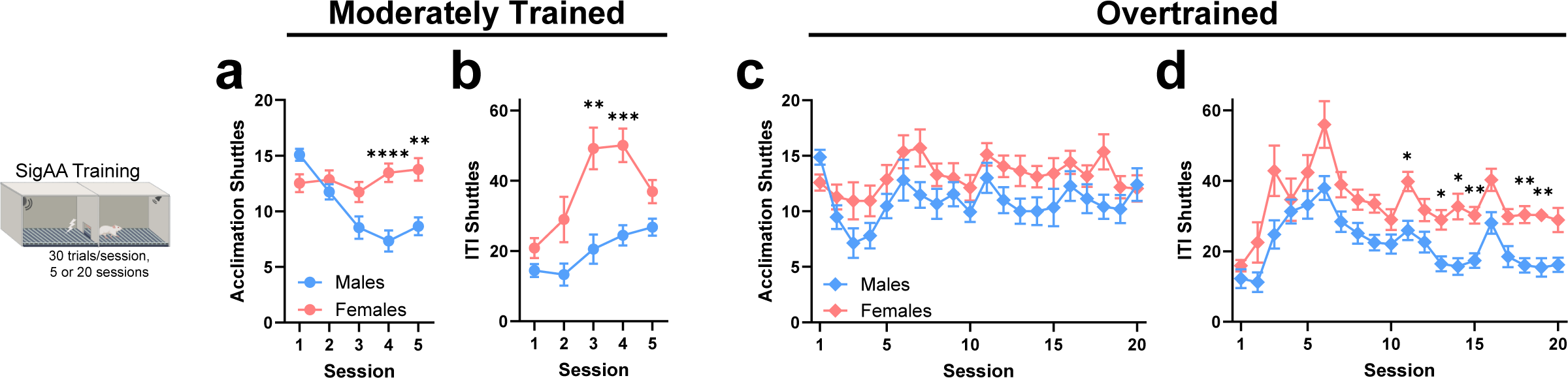
Female rats are slightly more active during SigAA training. Shuttles during acclimation and ITI periods for rats depicted in Fig 1. All data were analyzed by two-way ANOVAs followed by Šídák’s multiple comparisons tests. **a**, Acclimation shuttles for moderately trained male (n=35) and female (n=26) rats. Two-way ANOVA followed by Šídák’s test: F_4,236_=12.34. Group comparisons, Male versus Female, Session 4 ****p<0.0001, Session 5 **p= 0.0013. **b**, ITI shuttles for the same moderately trained rats. Two-way ANOVA followed by Šídák’s test: F_4,236_=3.76. Group comparisons, Male versus Female, Session 3 **p=0.0013, Session 4 ***p=0.0002. **c**, Acclimation shuttles for overtrained male (n=15) and female (n=17) rats. Two-way ANOVA: F_19,570_=1.652. **d**, ITI shuttles for the same overtrained rats. Two-way ANOVA followed by Šídák’s test: F_19,570_=0.8593 (Session x Sex), F_1,30_=20.56 (Sex). Group comparisons, Male versus Female, Session 11 *p=0.025, Session 13 *p= 0.025, Session 14 *p= 0.01, Session 15 **p= 0.006, Session 18 **p= 0.0019, Session 19 **p=0.0013. All data in the figure are shown as mean ± s.e.m.

**Extended Fig. 2.**
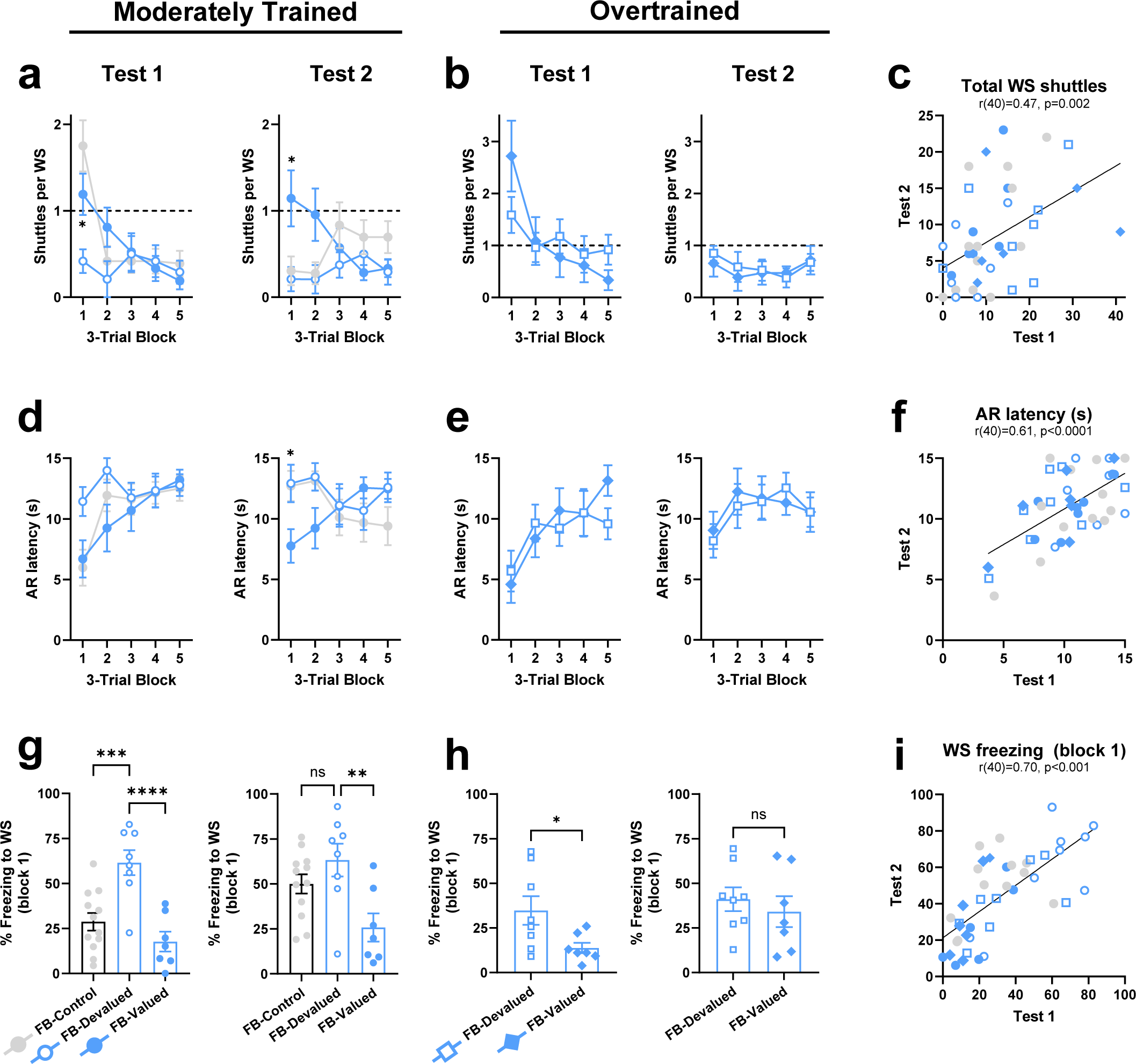
Devaluation test-retest reliability for male rats. We conducted two Devaluation Tests in anticipation of within-subject studies exploring the neural substrates of goal-directed vs. habitual avoidance using DREADDs. These studies require that each rat receive a test after injection with Vehicle or the DREADD ligand (counterbalanced). For this strategy to be effective, it is important that there is high test-retest reliability for our avoidance and freezing measures. Here we show the results of Test 1 and Test 2 for all moderately trained and overtrained rats in Fig. 2. Importantly, correlations between Test 1 and Test 2 behavior were very strong for all three measures indicating high test-retest reliability. **a**, Shuttles per WS for moderately trained rats depicted in Fig. 2 during Devaluations Tests 1 (left) and 2 (right). Two-way ANOVAs followed by Dunnett’s tests: Test 1, F_8,96_=4.38; Test 2: F_8,96_=4.595. Group comparisons, FB-Devalued versus FB-Valued, Test 1, Block 1 *p=0.034, Test 2, Block 1 *p=0.0495. **b**, Shuttles per WS for overtrained rats during Devaluation Tests 1 (left) and 2 (right). Two-way ANOVAs: Test 1, F_4,52_=3.02; Test 2, F_4,52_=0.3925. **c**, Pearson correlation of total WS shuttles (Test 1 vs Test 2) for all rats depicted in **a** & **b** (r_40_=0.4680, p=0.0018). **d**, Latency to first AR for moderately trained rats during Devaluation Tests 1 (left) and 2 (right). Two-way ANOVAs followed by Dunnett’s tests: Test 1, F_8,96_=1.82; Test 2, F_8,96_=2.85. Group comparisons, FB-Devalued versus FB-Valued, Test 2, Block1 *p=0.05. **e**, Latency to first AR for overtrained rats during Devaluation Tests 1 (left) and 2 (right). Two-way ANOVAs: Test 1, F_4,52_=1.43; Test 2, F_4,52_=0.35. **f**, Pearson correlation of average latency to first AR (Test 1 vs Test 2) for all rats depicted in **d** & **e** (r_40_=0.6063, p<0.0001). **g**, WS freezing during Block 1 for moderately trained rats in Devaluation Tests 1 (left) and 2 (right). One-way ANOVAs followed by Dunnett’s tests: Test 1, F_2,24_=13.87; Test 2, F_2,24_=5.85. Group comparisons, FB-Devalued versus FB-Control, Test 1 ***p=0.0006, FB-Devalued versus FB-Valued, Test 1 ****p<0.0001, Test 2 **p=0.0047. **h**, WS freezing during Block 1 for overtrained rats in Devaluation Tests 1 (left) and 2 (right). Unpaired t-tests: Test 1, t_13_=2.34 *p=0.036; Test 2, t_13_=0.65 p=0.52. **i**, Pearson correlation of average WS freezing (Block 1; Test 1 vs Test 2) for all rats depicted in **g** & **h** (r_40_=0.7031, p<0.0001). Data in panels **a**, **b**, **d**, **e**, **g** & **h** are shown as mean ± s.e.m.

**Extended Fig. 3.**
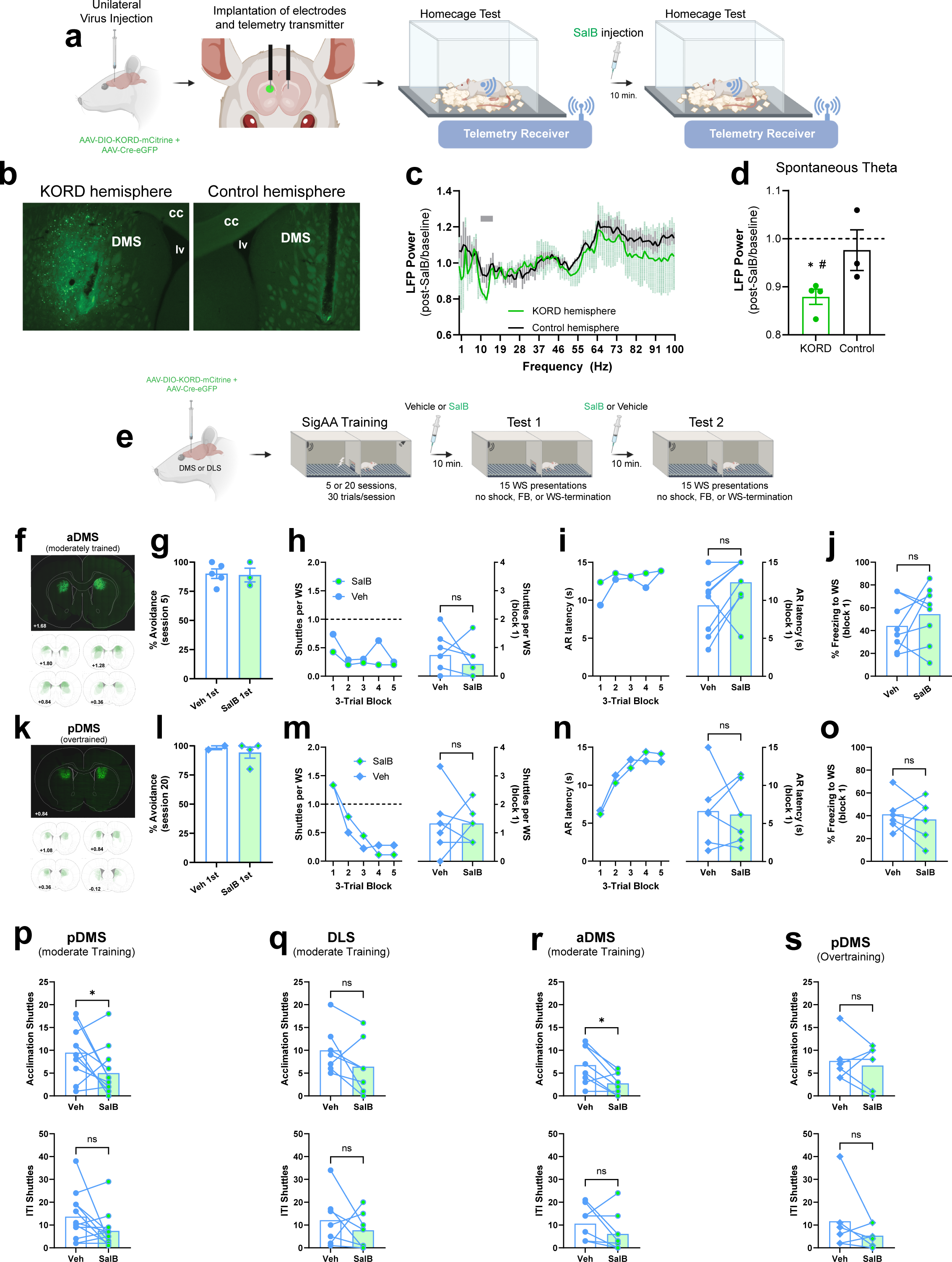
KORD validation, relevant null effects, and non-avoidance shuttling. **a**, Schematic of *in vivo* electrophysiology experiment to validate KORD/SalB inhibition in DMS. **b**, Example of unilateral KORD expression in DMS and bilateral electrode tracks. **c**, Spontaneous LFP power in KORD versus Control hemispheres post-SalB injection (normalized to baseline, n=4 rats). Gray Bar denotes theta band. **d**, Spontaneous theta band LFP power in KORD versus Control hemispheres post-SalB injection (normalized to baseline). Paired t-tests: baseline versus post-SalB theta in KORD hemisphere, t_3_=7.67 ^#^p<0.01; post-SalB theta in KORD versus Control hemisphere, t_2_=2.81 *p=0.05; baseline versus post-SalB theta in Control hemisphere, t_2_=0.56 p=0.32. **e**, Schematic of the experimental paradigm evaluating aDMS after moderate training (n=8) and pDMS after overtraining (n=6). **f**, Top, example of bilateral KORD expression in aDMS. Bottom, Semitransparent overlays illustrating KORD expression in aDMS for individuals across the A-P axis. A-P coordinates are in millimeters relative to Bregma. **g**, SigAA performance during session 5 of training for aDMS rats assigned to receive Vehicle versus SalB in Test 1. Unpaired t-test: t_6_=0.16 p=0.88. **h**, Left, shuttles per WS for Vehicle versus SalB Tests in aDMS rats. Two-way ANOVA: F_4,56_=0.75. Right, shuttles per WS for Block 1 of Tests. Paired t-test: t_7_=1.195 p=0.27. **i**, Left, latency to first AR for Vehicle versus SalB Tests in aDMS rats. Two-way ANOVA: F_4,56_=0.86. Right, latency to first AR during Block 1 of Tests. Paired t-test: t_7_=2.11 p=0.07. **j**, WS freezing for Block 1 of Vehicle versus SalB Tests in aDMS rats. Paired t-test: t_7_=1.08 p=0.32. **k**, Top, example of bilateral KORD expression in pDMS. Bottom, semitransparent overlays illustrating KORD expression in pDMS for individuals across the A-P axis. A-P coordinates are in millimeters relative to Bregma. **l**, SigAA performance during session 20 of training for pDMS rats assigned to receive Vehicle versus SalB in Test 1. Unpaired t-test: t_4_=0.58 p=0.60. **m**, Left, shuttles per WS for Vehicle versus SalB Tests in overtrained pDMS rats. Two-way ANOVA: F_4,40_=0.48. Right, shuttles per WS for Block 1 of Tests. Paired t-test: t_5_=0.00 p>0.99. **n**, Left, latency to first AR for Vehicle versus SalB Tests in overtrained pDMS rats. Two-way ANOVA: F_4,40_=0.415. Right, latency to first AR during Block 1 of Tests. Paired t-test: t_5_=0.24 p=0.82. **o**, WS freezing for Block 1 of Vehicle versus SalB Tests in overtrained pDMS rats. Paired t-test: t_5_=0.575 p=0.59. **p**, Top, Acclimation shuttles for Vehicle versus SalB Tests in moderately trained pDMS rats. Paired t-test: t_11_=2.305 *p=0.04. Bottom, ITI shuttles for the same rats. Paired t-test: t_11_=1.84 p=0.09. **q**, Acclimation shuttles for Vehicle versus SalB Tests in moderately trained DLS rats. Paired t-test: t_6_=1.77 p=0.13. Bottom, ITI shuttles for the same rats. Paired t-test: t_6_=0.84 p=0.43. **r**, Acclimation shuttles for Vehicle versus SalB Tests in moderately trained aDMS rats. Paired t-test: t_7_=3.35 *p=0.01. Bottom, ITI shuttles for the same rats. Paired t-test: t_7_=1.48 p=0.18. **s**, Acclimation shuttles for Vehicle versus SalB Tests in overtrained pDMS rats. Paired t-test: t_5_=0.51 p=0.63. Bottom, ITI shuttles for the same rats. Paired t-test: t_5_=1.26 p=0.265. cc: corpus callosum, lv: lateral ventricle. All data in the figure are shown as means. Error bars (s.e.m.) are shown for between-subject analyses.

**Extended Fig. 4.**
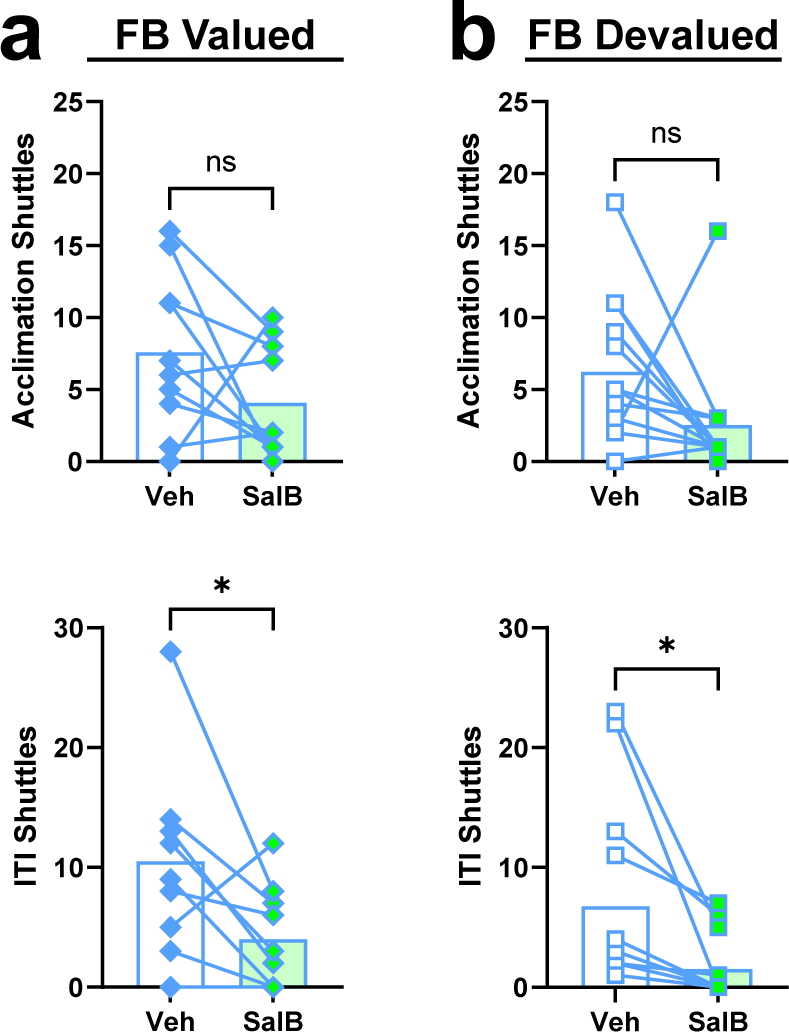
Acclimation and ITI shuttles for DLS habit experiment. **a**, Top, Acclimation shuttles for Vehicle versus SalB Tests in overtrained, FB-Valued rats depicted in Fig. 4. Paired t-test: t_9_=1.62 p=0.14. Bottom, ITI shuttles for the same rats. Paired t-test: t_9_=2.78 *p=0.02. **b**, Top, Acclimation shuttles for Vehicle versus SalB Tests in overtrained, FB-Devalued rats (n=13). Paired t-test: t_12_=1.87 p=0.09. Bottom, ITI shuttles for the same rats. Paired t-test: t_12_=2.76 *p=0.02. All data in the figure are shown as mean + individual scores.

**Extended Fig. 5.**
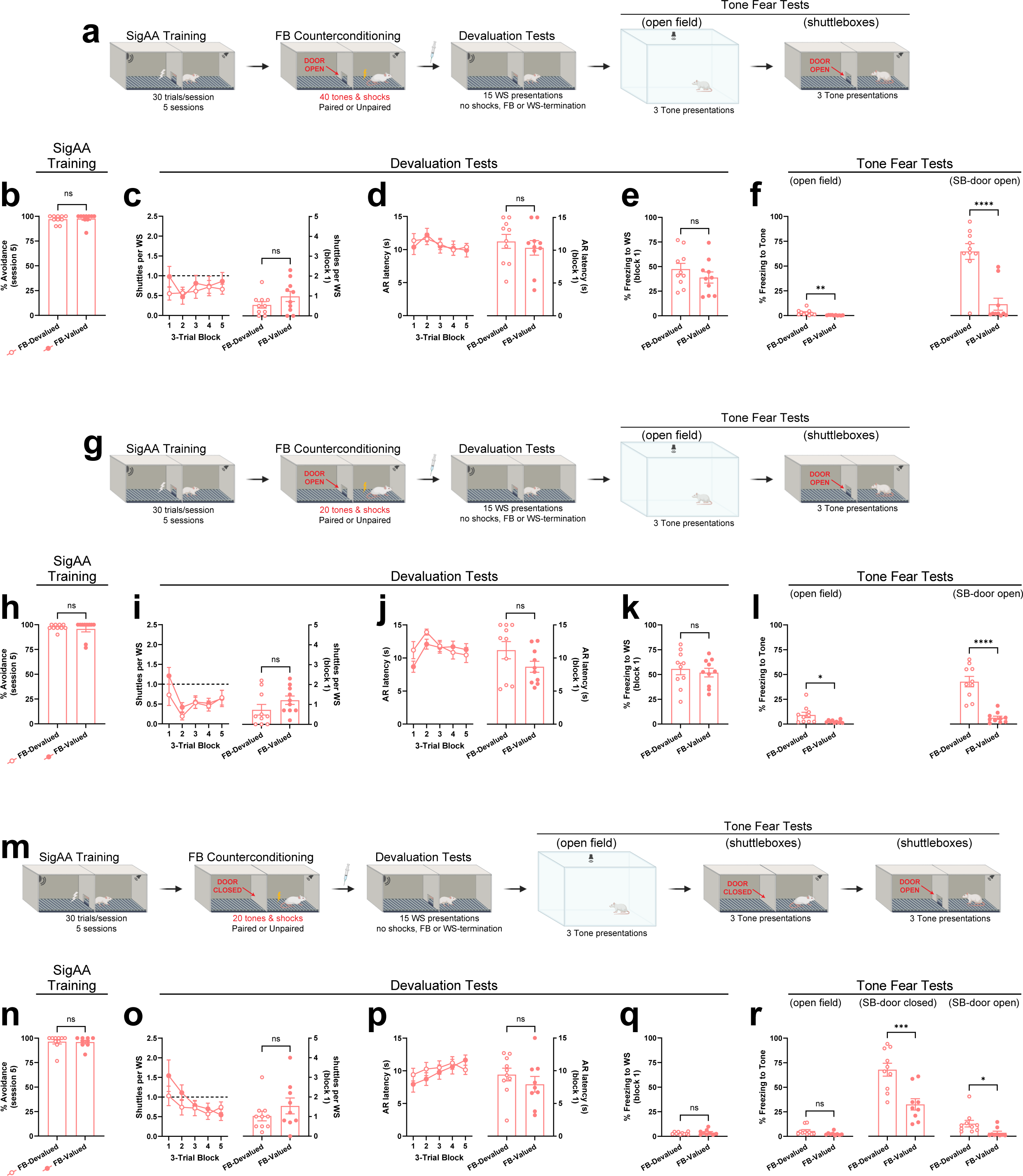
Devaluation of response-produced safety signals in the SigAA context (females). **a**, Schematic of the OPEN40 experimental protocol (OPEN refers to the shuttlebox door and 40 refers to the number of tones and shocks delivered during counterconditioning). Counterconditioning occurred in the shuttleboxes and was otherwise identical to previous experiments. **b**, SigAA performance during session 5 of training for rats assigned to FB-Devalued (n=10) and FB-Valued (n=10) groups. Unpaired t-test: t_18_=0.32 p=0.75. **c**, Left, shuttles per WS for both Devaluation Tests. Two-way ANOVA: F_4,72_=1.19. Right, shuttles per WS for Block 1 of Devaluation Tests. Unpaired t-test: t_18_=1.42 p=0.17. **d**, Left, latency to first AR for both Devaluation Tests. Two-way ANOVA: F_4,72_=0.41. Right, latency to first AR during Block 1 of Devaluation Tests. Unpaired t-test: t_18_=0.64 p=0.53. **e**, WS freezing for Block 1 of Devaluation Tests. Unpaired t-test: t_18_=1.06 p=0.30. **f**, Tone freezing in the novel open field (left) and shuttleboxes (door open; right). Unpaired t-tests: t_18_=2.997 **p=0.008 (left), t_18_=5.34 ****p<0.0001 (right). **g**, Schematic of the OPEN20 experimental protocol, which was identical to OPEN40 except only 20 tones & shocks were delivered during counterconditioning. **h**, SigAA performance during session 5 of training for rats assigned to FB-Devalued (n=10) and FB-Valued (n=10) groups. Unpaired t-test: t_18_=0.55 p=0.59. **i**, Left, shuttles per WS for both Devaluation Tests. Two-way ANOVA: F_4,72_=1.265. Right, shuttles per WS for Block 1 of Devaluation Tests. Unpaired t-test: t_18_=1.405 p=0.18. **j**, Left, latency to first AR for both Devaluation Tests. Two-way ANOVA: F_4,72_=2.25. Right, Average latency to first AR during Block 1 of Devaluation Tests. Unpaired t-test: t_18_=1.6 p=0.12. **k**, WS freezing for Block 1 of Devaluation Tests. Unpaired t-test: t_18_=0.51 p=0.62. **l**, Tone freezing in the novel open field (left) and shuttleboxes (door open; right). Unpaired t-tests: t_18_=2.42 *p=0.03 (left), t_18_=6.64 ****p<0.0001 (right). **m**, Schematic of the CLOSED20 experimental protocol, which was identical to OPEN20 except the door connecting shuttlebox chamber sides was closed during counterconditioning and an additional Tone Fear Test was conducted with the shuttlebox door closed. **n**, SigAA performance during session 5 of training for rats assigned to FB-Devalued (n=10) and FB-Valued (n=9) groups. Unpaired t-test: t_17_=0.14 p=0.89. **o**, Left, shuttles per WS for both Devaluation Tests. Two-way ANOVA: F_4,68_=1.49. WS shuttles for Block 1 of Devaluation Tests. Unpaired t-test: t_17_=1.11 p=0.28. **p**, Left, latency to first AR for both Devaluation Tests. Two-way ANOVA: F_4,68_=1.32. Right, latency to first AR during Block 1 of Devaluation Tests. Unpaired t-test: t_17_=0.96 p=0.35. **q**, WS freezing for Block 1 of Devaluation Tests. Unpaired t-test: t_17_=0.84 p=0.41. **r**, Tone freezing in the novel open field (left), shuttleboxes with door closed (middle), and shuttleboxes with door open (right). Unpaired t-tests: t_17_=1.79 p=0.09 (left), t_17_=4.00 ***p=0.0009 (middle), t_17_=2.38 *p=0.03 (right). All data in the figure are shown as mean ± s.e.m.

**Extended Fig. 6.**
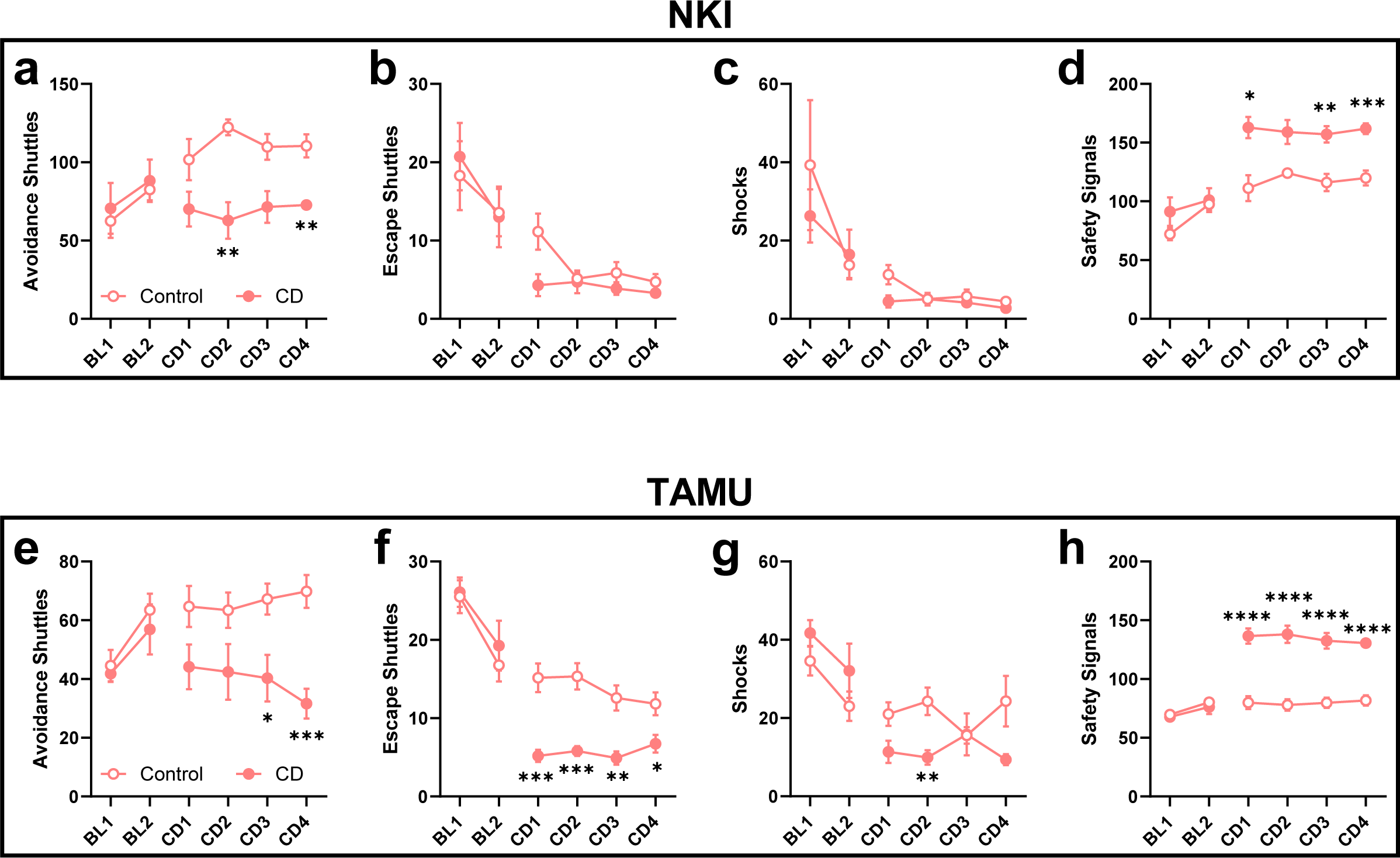
Replication of contingency degradation effect on goal-directed avoidance in females at separate institutions. Identical experiments were conducted in parallel at NKI (panels **a**-**d**) and TAMU (panels **e**-**h**) in females. Results are presented separately here but were combined for analysis in Fig. 6 given a similar pattern of results. Note that rats were matched into equal groups before the CD phase and there were no differences in baseline behavior (BL1-2) for any measure in either experiment. **a**, Right, CD rats (n=7) avoided less than Control rats (n=7) during the contingency degradation phase. Two-way ANOVA followed by Šídák’s test. F_3,36_=1.13 (Session x Group), F_1,12_=23.06 (Group). Group comparisons, CD versus Control, CD2 **p=0.006, CD4 **p=0.0060. **b**, Right, CD rats escaped less than Control rats during the contingency degradation phase. Two-way ANOVA: F_3,36_=2.71 (Session x Group), F_1,12_=6.00 *p=0.03 (Group). **c**, Right, CD and Control rats received similar patterns of shock during the contingency degradation phase. Two-way ANOVA: F_3,36_=2.36 (Session x Group), F_1,12_=4.13 (Group). **d**, Right, CD rats received more safety signals than Control rats during the contingency degradation phase. Two-way ANOVA followed by Šídák’s test: F_3,36_=0.48 (Session x Group), F_1,12_=37.68 (Group). Group comparisons, CD versus Control, CD1 *p=0.015, CD3 **p=0.007, CD4 ***p=0.0009. **e**, Right, CD rats (n=11) avoided less than Control rats (n=12) during the contingency degradation phase. Two-way ANOVA followed by Šídák’s test: F_3,63_=2.18 (Session x Group), F_1,21_=10.10 (Group). Group comparisons, CD verus Control, CD3 *p= 0.045, CD4 ***p=0.0002. **f**, Right, CD rats escaped less than Control rats during the contingency degradation phase. Two-way ANOVA followed by Šídák’s test: F_3,63_=3.423. Group comparisons, CD versus Control, CD1 ***p=0.0006, CD2 ***p=0.0005, CD3 **p=0.002, CD4 *p=0.045). **g**, Right, CD rats received fewer shocks than Control rats during the contingency degradation phase. Two-way ANOVA followed by Šídák’s tests: F_3,63_=3.12. Group comparisons, CD versus Control, CD2 **p=0.009. **h**, Right, CD rats received more safety signals than Control rats during the contingency degradation phase. Two-way ANOVA followed by Šídák’s test: F_3,63_=1.23 (Session x Group), F_1,21_=63.09. Group comparisons, CD versus Control, CD1 ****p<0.0001, CD2 ****p<0.0001, CD3 ****p<0.0001, ****p<0.0001. BL: baseline, CD: contingency degradation, TAMU: Texas A&M University, NKI: Nathan Kline Institute. All data in the figure are shown as mean ± s.e.m.

## Footnotes

[1] Although SalB is reported to be inert, we noticed effects of SalB injections in some of our male rats, including naïve rats not expressing KORD. Effects included a suppression of locomotor activity, squinting, and labored breathing that did not occur with vehicle injections. The effects appeared to subside within 10-20 minutes and could not account for the avoidance effects we observed with SalB/KORD. We did not observe similar effects in females who all received less total drug than males.

[2] Our results do not rule out additional roles for WS-termination and/or US-omission in the negative reinforcement of AA, especially early in training before FB cues develop inhibitory properties. However, conditioned inhibition likely develops faster than is currently appreciated. For instance, we have observed evidence for summation in as little as three trials with an explicitly unpaired fear conditioning procedure (Fig. S1 in reference^66^)

